# Ubiquitin-derived artificial binding proteins targeting oncofetal fibronectin reveal scaffold plasticity by β-strand slippage

**DOI:** 10.1101/2023.07.04.547672

**Authors:** Anja Katzschmann, Ulrich Haupts, Anja Reimann, Florian Settele, Manja Gloser-Bräunig, Erik Fiedler, Christoph Parthier

**Author notes:** **Corresponding authors** Christoph Parthier, Erik Fiedler.

## Abstract

Affilin proteins (artificial binding proteins based on the ubiquitin scaffold) were generated using directed protein evolution to yield *de-novo* variants that bind the extra-domain B (EDB) of oncofetal fibronectin, an abundant tumor marker in fetal and neoplastic tissues. Structures of two EDB-specific Affilin molecules reveal striking structural plasticity of the ubiquitin scaffold, characterized by β-strand slippage, leading to diverse register shifts of the β5 strands. This recruits residues from β5 to a loop region, enhancing the target-binding interface. The observed β-strand rearrangements, manifested by pressure of selection for target binding, challenge the accepted paradigm that directed evolution of binding proteins should base on rigid frameworks. Fold plasticity allowing β-strand slippages enhances the evolutionary potential of proteins beyond “simple” mutations significantly and provides a general mechanism to generate residue insertions/deletions in proteins. They are however difficult to predict, underlining the need for caution in interpretation of structure-activity relationships of evolved proteins.

## Introduction

The choice of a suitable, well-understood structural scaffold usually sets the course for the development, producibility and potential future applications of artificial binding proteins generated by directed evolution. Tailored for a broad spectrum of usage in medicine, biotechnology and research, an increasing variety of artificial binding proteins, derived from non-immunoglobulin-based scaffolds, have been established, featuring favorable biophysical properties and potential to specifically bind any target molecule ^1, 2^. Most of these scaffolds employ rather compact, naturally occurring, stable and structurally well-characterized protein modules, comprising secondary structural elements such as a-helices, β-sheets and loops to varying degrees. They include lipocalins ^3^, ankryrin repeat proteins ^4^, staphylococcal protein A ^5^, fibronectin ^6^, g-crystallin ^7^, ubiquitin ^8, 9^ and others ^10^.

Typically, these modules are chosen as structural platforms or building blocks for the directed evolution of *de-novo* binding properties, the latter being achieved by specific amino acid exchanges at selected positions. Screening of combinatorial scaffold protein libraries using appropriate selection techniques allows subsequent isolation of candidate binders with the desired properties. In spite of the power of combinatorial evolution, the success of the approach is strongly dependent on the selection of the amino acid positions to be diversified. This process usually relies on structural knowledge and considers the overall fold of the scaffold protein to be essentially fixed, i.e. the general content and spatial arrangement of secondary structure elements ^11, 12^.

The structure of ubiquitin (Ub), a 76 amino acid signalling protein conserved among all eukaryotes, features a single five-stranded β-sheet enclosing a short α-helix. Approaches to adopt a single Ub chain as a scaffold for the generation of binding proteins ^8, 13^ were extended by the development of diubiquitin-based scaffolds (Affilin molecules), generated by genetic fusion of two Ub domains to target a tumor-specific fibronectin (Fn) variant, oncofetal Fn ^14^. The alternative splice variant of Fn comprises the extra-domain B (EDB), specifically expressed in tissues of tumor-associated angiogenesis and other neoplastic processes ^15, 16^.

Here we report how directed evolution of Affilin molecules targeting oncofetal Fn results in unusual structural rearrangements by different β-strand shifts, reflecting a striking plasticity of the diubiquitin scaffold and enabling the formation of the target binding site. We also show the potential of these binding proteins by addressing EDB-expressing cells with a chimeric Afilin-IL2-fusion as targeting modules in tumor-directed diagnostics and therapy.

## Results

### Generation of Affilin molecules targeting EDB of oncofetal fibronectin

The development of the dibiquitin-based Affilin Af1 with evolved binding properties against human oncofetal Fn has been previously described ^14^. Following a related approach, we employed the diubiquitin scaffold, to generate the Affilin Af2 using a modified library setup, an improved selection procedure and an altered affinity maturation process (supplementary discussion). In brief, two libraries of monomeric Ubiquitin (variant Ub-WAA) were randomized at 7 positions (residues 6 and 8, located in strand β1/loop β1β2, 62-66, located in loop α2β5/strand β5, figure 1e, supplementary figure S1) and subsequently linked to yield a library of diubiquitin variants, constituting the basis for selection of EDB-binding Affilin molecules by phage display (PD). The selected candidate, variant Af2p, comprised 14 evolved amino acid changes versus diUb-WAA: Lys6His, Leu8Asp, Gln62Asp, Lys63Pro, Glu64Gln, Ser65Leu, Thr66Lys, Lys85Thr, Leu87Gln, Lys141Asp, Glu142Tyr, Ser143Arg, Thr144Tyr and the deletion ΔGln140. In binding analysis, Af2p displayed nanomolar affinities to the target 67B89 and no binding to the Fn fragment 6789 lacking the EDB, while thermal stability decreased significantly (ΔT_m_=-18K,) compared to the starting variant diUb-WAA (table 1, extended data figures 1 and 4). During subsequent affinity maturation, error-prone PCR was used to introduce amino acid exchanges at any position, followed by stringent ribosome display (RD) selection of molecules binding to the target 67B89. Selection pressure was increased over four rounds of RD to select for binders with a low off-rate. Target binding affinity and specificity, thermal stability and recombinant producibility were assessed to nominate the final Affilin variant Af2s differing from Af2p by two additional mutations (Pro38Gln and Tyr143Phe) and a deletion (ΔIle78) in the linker region between the N- and C-terminal Ub domains. The maturation process yielded higher binding affinities for Af2s (and streptag-free Af2) as well as a significant recovery of thermal stability (ΔT_m_=+9K versus Af2p) (supplementary discussion, figure 1a and 1b, table 1, extended data figures 1 and 4).

**Table 1.**
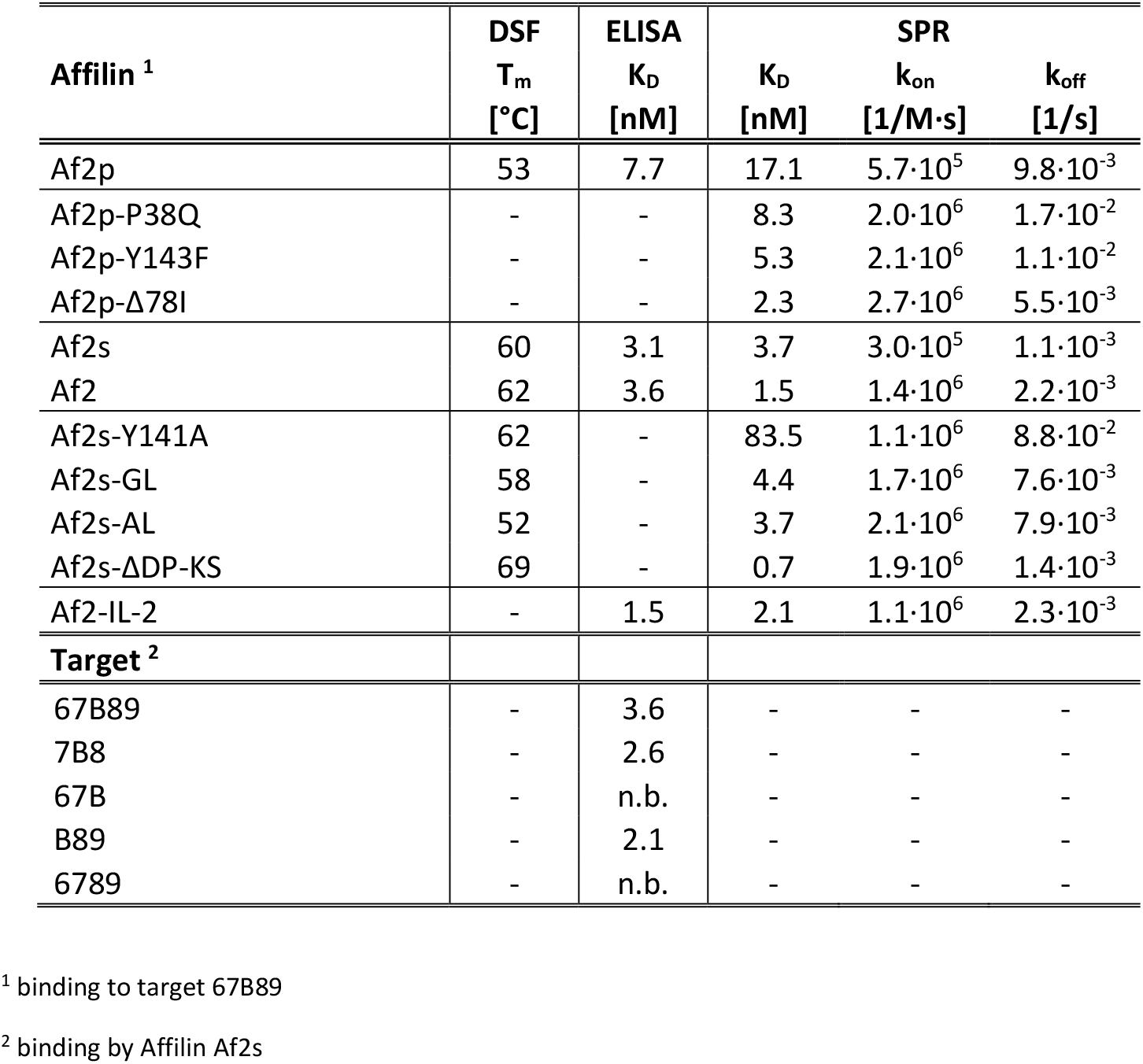

**Figure 1.**
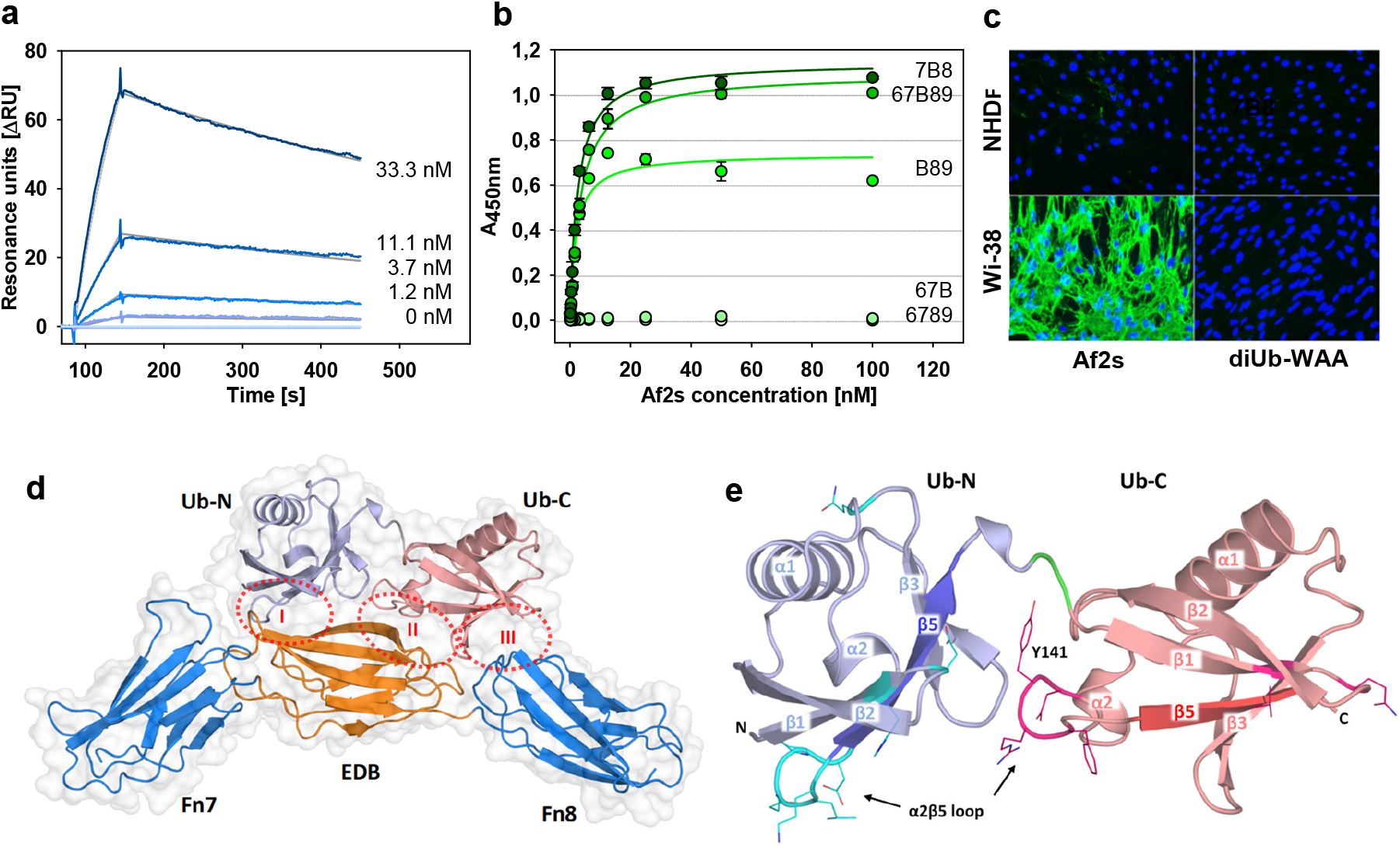
Binding analysis and structure of Affilin Af2 targeting onocfetal fibronectin (binding parameters are given in table1). (**a**) SPR analysis of Affilin variant Af2s binding to the immobilized target 67B89 at different concentrations of Af2s. (**b**) Af2s binding to different target variants comprising EDB, analysed by ELISA. (**c**) EDB-specific binding of Af2s to human fetal lung fibroblast cells (Wi-38) detected by immunofluorescence (green), cell nuclei stained with DAPI (blue). (**d**) Crystal structure of the Af2:7B8 complex revealing three binding regions (I-III, dotted red elipses). **(e)** Overall structure of 7B8-bound Af2 highlighting the secondary structure elements with β5-strands of Ub-N (dark blue) and Ub-C (dark red), evolved residues depicted as lines, colored by atom type (C-atoms of Ub-N cyan, C-atoms of Ub-C magenta, N atoms blue, O atoms red). Linker residues colored in green.

As crystallization of Af2 in complex with 67B89 was unsuccessful, truncated versions of the target were generated (supplementary figure S2a) where variants 7B8 and B89 exhibited very similar affinities to Af2s as 67B89, while no binding was observed for 67B and the control 6789, lacking EDB (table 1, figure 1b). This suggests EDB and Fn domain 8 (Fn8) predominantly contribute to Affilin binding.

Af2s preserves specific binding to oncofetal fibronectin in cell culture: using an immunofluorescence assay, Af2s showed EDB-specific binding to human fetal lung fibroblast Wi-38 cells, expressing a high level of oncofetal fibronectin, In contrast, binding to neonatal human dermal fibroblast cells (NHDF), a cell line with low EDB expression, was significantly reduced (figure 1c). Diubiquitin (variant diUb-WAA) showed no detectable binding and confirmed the targeting of EDB-expressing cells is mediated by the evolved binding properties of the Affilin.

### Target-bound Af2 reveals distinct register shifts in the β5 strands

HPLC-SEC analysis of the purified Af2:7B8 complex indicated a single species with an apparent molecular mass of about 45 kDa, corresponding to a 1:1 complex (supplementary figure S2b). Crystals of the Af2:7B8 complex, diffracting to 2.3Å, comprised three complexes in the asymmetric unit with slight variations between the individual molecules (figure 1d, supplementary figure S3, supplementary discussion). The crystal structure demonstrates the modular architecture of the Affilin, consisting of an N-terminal Ub-like domain (Ub-N) tethered to a C-terminal Ub-like domain (Ub-C) by a two-amino-acid linker (figure 1e). The bound target 7B8 adopts an extended conformation of three head-to-tail-linked Fn domains (Fn7, EDB, Fn8) with Fn7 and Fn8 in slightly kinked orientation relative to the central EDB. The structure confirms that only EDB and Fn8 interact with Af2, while Fn7 is not directly involved. The two Ub-like domains of Af2 together form a total binding interface of 1179 Å^2^, with the major contribution from Ub-C (811 Å^2^) versus Ub-N (368 Å^2^) (supplementary figure S9a-c). Three sub-interfaces (I-III, figure 1d) constitute the structural basis of target recognition by Af2. Interface I between Ub-N and EDB includes evolved residues from the α2β5 loop and strand β1, but also original (non-evolved) amino acids from β1 and β5 of Af2, which interact with EDB residues located in β-strands C/C’ and loop regions BC/C’E (figure 2a). Interface II is formed between Ub-C and EDB, comprising evolved residues from the α2β5 loop and a non-evolved R149 from β5 of Ub-C that interacts with EDB residues from β-strand C’ and the EF loop (figure 2b). Interface III involves interactions between Fn8 and Ub-C and is devoid of any evolved residues. Ub-wt residues (conserved in Af2) located in strands β3/β5 and the β3β4-loop of Ub-C, interact with amino acids of the FG loop of Fn8 (extended data figure 2). A detailed description of key intermolecular interactions underlying 7B8 binding is given in the supplementary discussion. Similar modes of 7B8 recognition have been described for Anticalins (lipocalin-derived artificial binding proteins) evolved against oncofetal fibronectin ^17, 18^. In spite of their distinct binding interfaces, they bind 7B8 in a surprisingly similar target conformation via EDB and the Fn8 domain, as observed for Affilin Af2 (supplementary figure S5).

**Figure 2.**
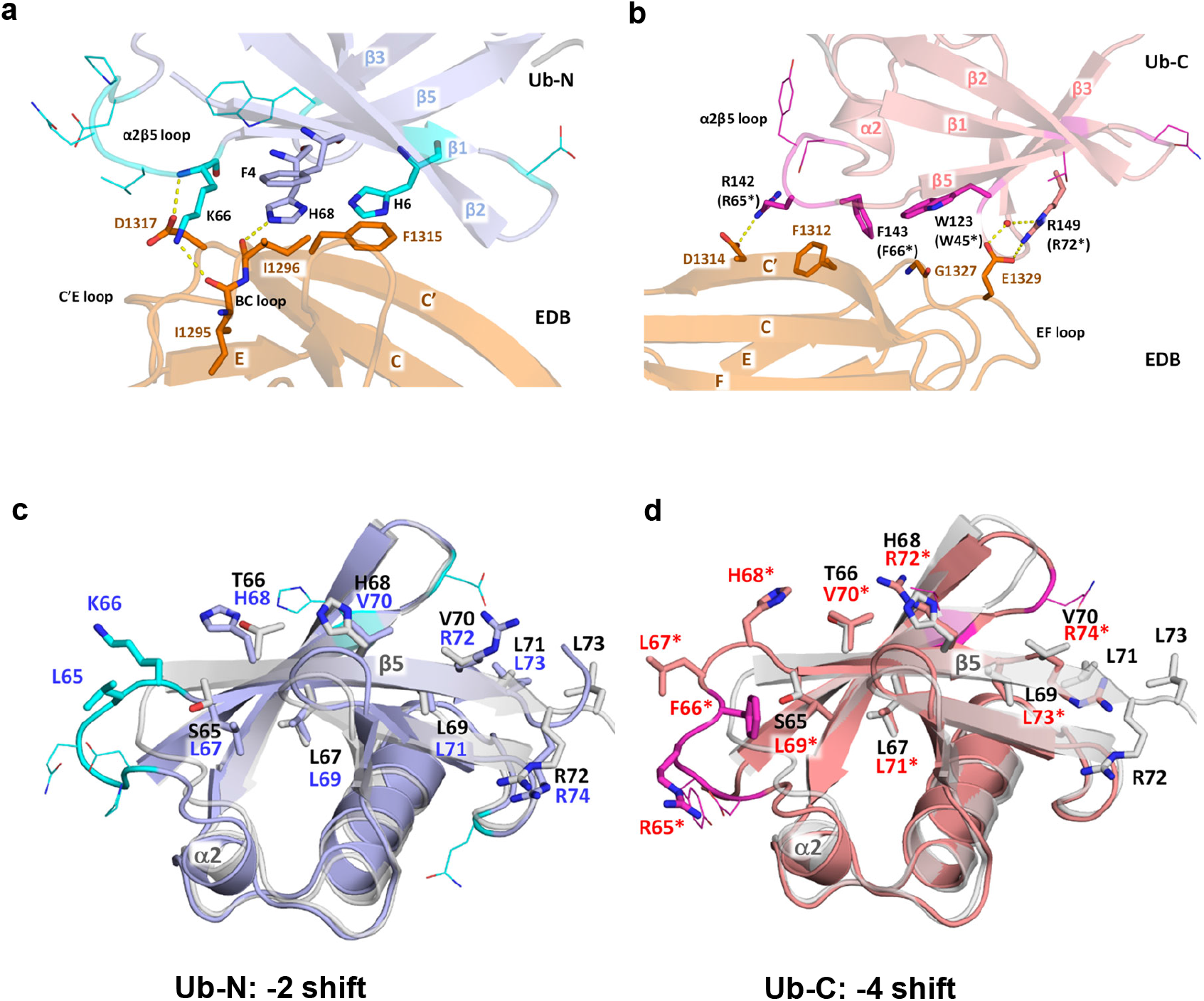
Molecular interactions contributing to EDB binding by Af2 exhibiting -2 and -4 register shifts in β5. Residues coloring of C-atoms in cyan/light blue (Ub-N), magenta/light red (Ub-C), orange (EDB), white (Ub-wt), N-atoms in blue and O-atoms in red. (**a**) Detailed view of key interaction between Ub-N and EDB (interface I). Residues involved in binding shown as sticks, evolved amino acids of Ub-N depicted as lines, hydrogen bonds shown as yellow dashed lines. (**b**) Key interaction between Ub-C and EDB (interface II). Residues involved in binding shown as sticks, evolved amino acids of Ub-C depicted as lines, hydrogen bonds shown as yellow dashed lines. **(c)** Structural superposition of the Ub-N domain of Af2 (light blue, blue labels) displaying a -2 register-shift in β5 vs. Ub-wt (PDB id 1UBQ, white, black labels). Residues of β5 shown as sticks. Evolved Af2 residues shown in cyan. (**d**) Structural superposition of the Ub-C domain of Af2 (light red, red labels with asterisk, corresponding to residue numbering of Ub-N domain) revealing a -4 register-shift in β5 vs. Ub-wt (colored in white, black labels). Residues of β5 shown as sticks. Evolved Af2 residues shown in magenta.

A contribution of evolved scaffold residues to target binding is generally expected, being a direct result of the evolutionary process. In the 7B8:Af2 complex, most of these are located in the α2β5 loops of Ub-N and Ub-C (figure 1e). In contrast, the participation of original Ub-wt residues, conserved also in Af2 and located in b5 (e.g. H68, V147, R149), appear at first glance to be coincidental. Careful inspection of the structural differences between Af2 and Ub-wt ^19^ however revealed these and other residues of β5 occupy different positions in the Affilin structures (figures 2c and d). Distinct β-strand slippages at β5 have occurred in Ub-N and Ub-C of Af2, resulting in register shifts by -2 residues (Ub-N) or -4 residues (Ub-C), as unambiguously seen in the 2F_o_-F_c_ electron density of the Af2:7B8 complex (supplementary figure S6a-b). The register shifts are observed in all three complexes in the asymmetric unit. The shifts recruit two residues in Ub-N (L65/K66) and four amino acids in Ub-C (R142/R65*, F143/F66*, L144/L67*, H145/H68*, the asterisk designating the residue numbering corresponding to Ub-N) from β5 to the preceding α2β5 loop and relocate all subsequent residues of the Ub-domain in the same direction. This remarkable rearrangement appears to be due to the sequence repeat of four leucines (L67, L69, L71, L73) in β5 of both Ub domains, allowing the occupation of two hydrophobic grooves in the Ub domain core by leucines also present in the -2 or -4 register-shift of β5 (L67/L69 in Ub-wt, L69/L71 in Ub-N, L71*/L73* in Ub-C, supplementary figures S6c-d). Interestingly, amino acid deletions within the α2β5 loop, effectively shortening the loop extensions while preserving the register shift, can reverse the thermodynamic destabilization observed for the register-shifted variants (supplementary discussion).

While all Af2 residues recruited from b5 to the α2β5 loop are directly involved in target binding, other evolved residues contribute indirectly, with a peculiar role of Y141 (Y64*), also located in the α2β5 loop of Ub-C. With its sidechain packed in the interface between Ub-N and Ubi-C, it mediates critical domain contacts that apparently stabilize the domains in an orientation, optimal for target binding (figure 1e). The “locked” position of Y141 restricts also the conformation of the whole α2β5 loop in Ub-C, and thereby affects target binding. Substitution of Y141 with alanine (variant Af2s-Y141A) results in a stable Affilin variant that exhibits a drastically reduced binding affinity towards the target 67B89, compared to Af2s (table 1, extended data figure 3d). Accordingly, reduction of linker length between Ub-N and Ub-C (evolved deletion ΔIle78) contributed significantly to the gain of affinity during the maturation of Af2 (table 1).

### The structure of Af1 reveals β5 register-shifts in absence of the target

Attempts to crystallize the purified Af1:7B8 complex resulted in crystals of Af1 alone that diffracted to 2.2 Å, allowing elucidation of the structure of Af1 in the unbound state (supplementary table S1). Superposition of Af1 with the Af2:7B8 complex reveals different orientations of the two Ub-like domains: Ub-C of Af1 is rotated about 180° with respect to Ub-N about an axis almost parallel to the linker (extended data figure 5). In free Af1, the evolved residue W142 (W63*) from the α2β5 loop of Ub-C is solvent-exposed. This is in contrast to the corresponding Af2 residue Y141 (Y64*) in the complex, which mediates interdomain contacts between Ub-N and Ub-C. Despite distinct mutations evolved for Ub-N and Ub-C of Af1, the two domains adopt nearly identical overall structures (supplementary figure S8a/b). Interestingly, both Af1 domains exhibit a -2 register shift of β5, even in the absence of a bound target. As observed for Af2 Ub-N, the shifts in Af1 recruit the evolved residues at positions 65/66 (65*/66*) from β5 to the α2β5 loop, relocating the subsequent amino acids in the N-terminal direction (supplementary discussion). The structures suggest that the -2 register-shifted states of Af1 and Af2 have evolved to bind the target oncofetal fibronectin. Nevertheless, as several evolved residues differ between Af1 and Af2 (supplementary figure S1), the precise binding modes of the two Affilins are expected to be different. Clearly, the -2 register shift in Af1 Ub-C is not compatible with the binding mode observed in the Af2:7B8 complex (comprising a -4 shift in Ub-C) and would result in clashes of Af1 Ub-C with the EDB and Fn8 domains. In contrast, the Ub-N domains of Af1 and Af2 share the same register shift and could allow similar interactions between Af1 Ub-N and EDB (supplementary figure S8c). Three different binding modes for Af1 are conceivable: (i) Af1 binds the target completely differently to Af2; (ii) Af1 Ub-N interacts with the target as in the Af2:7B8 complex, Af1 Ub-C recognizes a different epitope of oncofetal fibronectin; or (iii) Ub-N and Ub-C of Af1 bind the target in a manner similar to the Af2:7B8 complex, which would require additional conformational changes in Ub-C. In the last scenario, Af1 Ub-C would have to undergo a significant domain reorientation to contact the target protein, as well as a binding-induced extension of the register shift from -2 to -4, recruiting Af1 residues into positions suitable for target binding.

### Chimeric Affilin-Interleukin-2 - a potential anti-tumoral agent targeting oncofetal fibronectin

Based on the specific EDB-binding properties of the Affilin, its potential application as a targeting unit was investigated. In conceptual analogy to immunocytokines, human IL-2 was genetically fused as a “payload” to Af2. Targeted delivery of the cytokine IL-2, to e.g. sites of tumoral angiogenesis, can augment cellular cytotoxicity by local T-cell stimulation ^20, 21^. As chimeric Af2-IL-2 expressed in *E*.*coli* resulted in the formation of insoluble inclusion bodies, *in vitro* refolding of the fusion protein followed by standard purification procedures was employed to facilitate protein production. Functionality of the Affilin moiety of the chimeric protein was assessed by analysing the binding properties towards the target 67B89, revealing affinities of Af2-IL-2 in the low nanomolar range with binding kinetics very similar to Af2. Fusion of Af2 to IL-2 preserved specificity of the chimeric protein towards the target 67B89 with no binding to the off-target 6789 observed (table 1, figure 3a, extended data figure 4d). The Af2-IL-2 fusion binds specifically to Wi-38 cells (human fetal lung fibroblasts expressing a high level of oncofetal fibronectin) as shown by an immunofluorescence assay (figure 3b). In contrast, binding to a cell line with low EDB expression (NHDF) was negligible, demonstrating the chimeric Affilin-cytokine fusion is capable of targeting EDB-expressing cells *in vitro*. Cytokine activity was assayed using a cell line of IL-2-dependent murine cytotoxic T-cells (CTLL-2) with recombinant human IL-2 as a reference. Comparison of mean EC_50_ values revealed similar potencies of refolded Af2-IL-2 (EC_50_= 33 ± 15 pM) and recombinant IL-2 (EC_50_=41 ± 11 pM), respectively (figure 3c), indicating that both proteins, Affilin and cytokine, remained active in context of their genetic fusion, and that the Affilin does not significantly affect the anti-tumoral activity of the payload IL-2.

**Figure 3:**
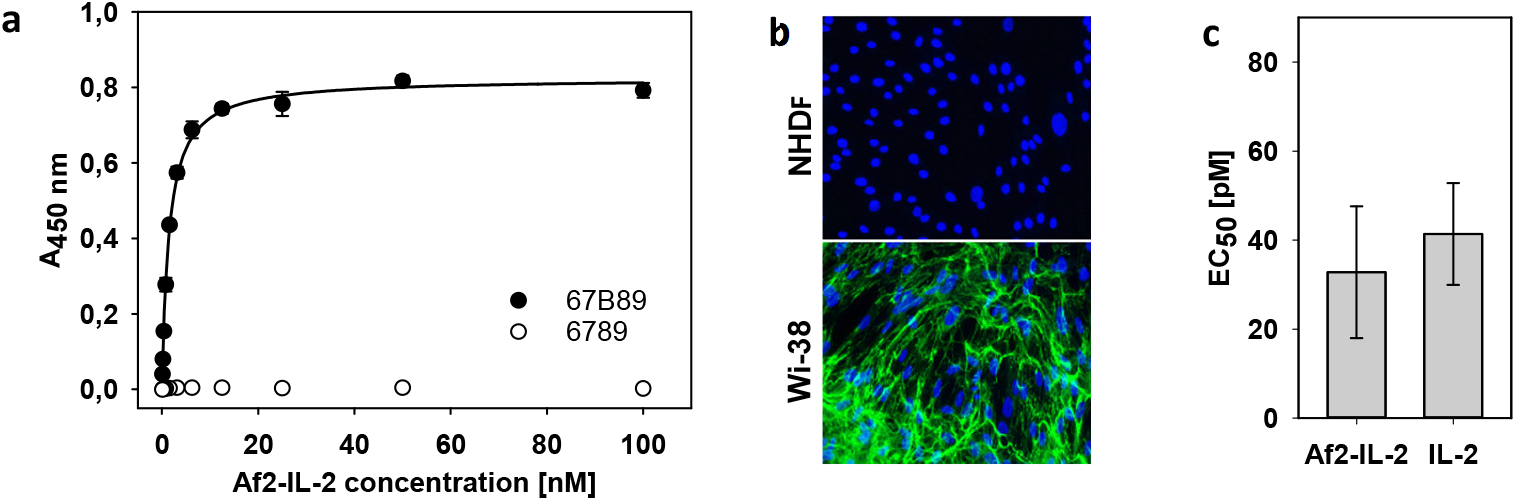
Binding and functional analysis of an EDB-specific Affilin-IL-2 chimera (binding parameters are given in table1). (**a**) Binding of Af2-IL-2 to the target 67B89 (filled circles) and lack of binding to the off-target 6789 (open circles) analyzed by ELISA. (**b**) EDB-specific targeting of Af2-IL2 to human fetal lung fibroblast cells (Wi-38) detected by immunofluorescence (green), cell nuclei stained with DAPI (blue). (**c**) IL-2-dependent growth of murine cytotoxic T-cells (CTLL-2) after application of Af2-IL-2, using recombinant human IL-2 as reference. IL-2 activity (given as EC_50_ values) from triplicate measurements was derived from the number of viable cells.

## Discussion

The directed evolution of proteins with engineered binding properties has employed many non-antibody scaffolds successfully, circumnavigating the structural complexity of immunoglobulins and derived molecules ^12^. For the most part, these artificial binding proteins have adopted the natural paradigm of antibody diversity, where structural adaptations to target binding are achieved predominantly via variable (evolved) residues framed in a rigid scaffold. Confirmed in numerous structural studies, this principle governs scaffold selection as well as library design ^11^. With the crystal structures of two independently evolved Affilin molecules Af1 and Af2 raised against the target oncofetal fibronectin, we have determined the structural basis of target recognition by a diubiquitin-based artificial binding protein, revealing a striking example of scaffold plasticity. Contrary to the structurally inert scaffold model, we observed variations of the central β-sheet register that result from apparent strand slippages of 2 or 4 residues. Register shifts were found for the β5-strand in each Ub domain of unbound Af1 (−2 shifts) and target-bound Af2 (−2 shift in Ub-N, -4 shift in Ub-C), respectively. The structural consequences are two-fold: i) each α2β5 loop is extended by two (−2 shift) or three residues (−4 shift, coinciding with the evolved deletion DQ140), and ii) all subsequent domain residues become relocated, recruiting non-evolved residues to positions where they now can contribute to target binding (figure 2, supplementary figure S1). Obviously, amino acid composition at the randomized positions as well as changes in scaffold structure have co-evolved in the selection process in order to acquire the desired binding properties.

Structural evidence for β-strand slippage in natural proteins is sparse, as it can be easily overlooked in sequence alignments and (low resolution) structures, impeding also their prediction. As such, they could be more common than current structural knowledge reflects. Wild-type ubiquitin is a prototype example, where it has been shown that β5 strand slippage is a natural feature of ubiquitin ^22^. During mitophagy (the clearance of damaged mitochondria), Ub-wt is phosphorylated at S65 by the Ub kinase PINK1 ^23^. This requires a transient -2 register shift of the Ub β5 strand to relocate S65 to an exposed position in the α2β5 loop, allowing PINK1 access to the substrate residue. In the absence of phosphorylation, the -2 register-shifted state of Ub-wt is sparsely populated (< 1%), whereas phosphorylation by PINK1 or mutations of Ub residues in β5 (e.g. T66V and L67N, variant Ub-TVLN) shift the conformational equilibrium significantly towards the register-shifted state ^22, 24-26^ (supplementary discussion, supplementary figures S9 d, e, h). A ubiquitin register shift of more than two residues, as found in Ub-C of Af2, has been reported only once before for an evolved ubiquitin variant (Ubv-G08) possessing 7 mutations spread over the strands β1, β4, β5 and the α2β5 loop. The crystal structure of Ubv-G08 in complex with its target revealed a -4 register shift of the β5 strand, with concomitant extension of the α2β5 loop, with both regions extensively involved in ligand binding ^27^ (supplementary discussion, supplementary figures S1 and S9 f, g, i).

Slippage of β-strands has also been described for a number of other proteins. GTP-binding proteins link strand slippage to GTP hydrolysis affecting membrane remodelling and filament formation ^28, 29^. The zymogen coagulation factor VII undergoes a three residue register shift during activation ^30^, bacterial flavin-dependent BLUF photoreceptors employ β strand slippage in the transition between dark and light state^31^ and in the *Vibrio cholerae* toxin HigB2 a single-residue β-strand register shift shuts off its mRNAse activity by flipping a catalytically important residue out of the active site ^32^.

β-strand register shifts represent one facet of a more general plasticity of β-sheets. The rearrangement of β sheets in donor strand complementation, both transient and permanent, reported in processes such as bacterial pilus formation ^33^, protease inhibition by serpins ^34^, ubiquitin ligase assembly ^35^ and chaperone activity ^36^, involve a similar reorganization of hydrogen bonding and side chain packing to that required for β-strand slippage. This applies also to domain-swapped proteins often formed through wholesale exchange of β-strands between molecules ^37^.

The β-strand slippage presented here provides a possible conceptual framework for the evolution of sequence insertions and deletions at the protein level to form neofunctional stable domains adding a fascinating aspect to (directed) protein evolution. Beyond “simple” mutations, register shifts may not only place evolved residues into new positions, they also affect a range of subsequent amino acids in the protein, recruiting them into a new environment. This multiplies the structural repertoire a scaffold protein can adopt, with the potential for optimizing (or creating *de novo*) binding sites. Although not all possible scaffold variations will result in stable structures, the power of the combinatorial approach, combined with an appropriate selection strategy may generate suitable candidates. The high-affinity oncofetal Fn-specific Affilins presented here demonstrate potential caveats in the interpretation of results from directed evolution and underline the importance of structural analyses for further development. The evolved binding proteins can be combined with a variety of biological functions (as shown for the chimeric Affilin-IL2 fusion), opening perspectives for diverse medical and biotechnological applications.

## Methods

### Production of fibronectin fragments

The genes for the target proteins, human fibronectin fragments containing EDB (67B89, 67B, 7B8 and B89, Uniprot ID P02751, isoform 7, variant C1232S) and off-target fragment 6789 lacking EDB (supplementary figure S2a) were obtained via gene synthesis (Geneart, Regensburg, Germany) and cloned into pET28a expression vector. The vectors were transferred into *E. coli* BL21 (Lucigen, Middleton, WI, USA). After protein expression cells were lysed and proteins purified via subsequent anionic exchange (Q-Sepharose FF XK 26/20 column), ammonium sulfate precipitation, hydrophobic interaction chromatography (HiTrap Phenyl HP column) and size exclusion chromatography (Superdex 200 XK 26/60 column). All chromatographic steps were carried out on an Aekta Explorer system (GE Healthcare, Freiburg, Germany). To allow selection of binders by ribosome and phage display, preferential N-terminal biotin labelling of the target 67B89 was achieved after dialysis of the sample against 50 mM sodium phosphate buffer pH 6.5 and subsequent incubation with a 30-fold molar excess of EZ-Link Sulfo-NHS-LC-Biotin reagent (Pierce, Rockford, IL, USA) for 24 hours at 4 °C. Non-coupled reagent was removed by dialysis against PBS pH 7.4.

### Affilin library construction

The Affilin scaffold used in this work consists of a linear fusion of two ubiquitin molecules (Uniprot ID P0CG47, residues 1-76). Prior to library generation three point mutations were introduced to improve spectrophotometric sensitivity ^38^ and manufacturability (F45W, G75A, G76A, yielding ubiquitin variant Ub-WAA). *In silico* analysis of protein stability effects exerted by mutation of candidate surface-exposed residues of ubiquitin identified 9 amino acid positions (2, 4, 6, 8, 62-66) which were selected for randomization in Affilin Af1, as described previously ^14^. For Affilin Af2, two library modules of monomeric Ub-WAA incorporating random amino acids (except cysteine) at positions 6, 8 and 62-66 were synthesized, differing only in codon usage (Morphosys, Martinsried, Germany). Introduction of *Mfe*I and *EcoR*I restriction sites via PCR, followed by digestion and ligation of the amplified fragments yielded a library of di-ubiquitin (diUb-WAA), resulting from the linear fusion of both Ub-WAA monomer libraries separated by a Gly-Ile-Gly linker. The diubiquitin library was further amplified via PCR using biotinylated primers to insert *Bsa*I restriction sites. Following digestion with *Bsa*I the insert was purified using magnetic streptavidin coupled beads (M-270 Dynabeads, Life Technologies, Carlsbad, CA) and subsequently ligated to phagemid pCD12, a derivative of phagemid pCD87SA ^39^. *E. coli* ER2738 cells (Lucigen, Middleton, WI) were transformed with the resulting phagemids by electroporation followed by single-colony PCR and DNA sequencing to assess correct size and sequence of the inserts. All transformed clones were purified using the QIAfilter Plasmid Maxi Kit (Qiagen, Hilden, Germany) to obtain the phagemid library.

### Phage display selection and screening of EDB-specific Affilin variants

Generation of Af1 is described elsewhere ^14, 40^. For Af2 precursors, Tat-mediated phage display (PD) ^39^ selection and screening of was performed at 20°C in a similar manner. Briefly, N-terminally biotinylated 67B89 target protein was immobilized on Streptavidin Dynabeads M-270 (Invitrogen, Carlsbad, CA, USA). The target-coated beads were blocked with BSA and incubated with a suspension of 3.4·10^12^ phages in the presence of a 10-fold molar excess of the off-target (variant 6789), followed by washing the beads with PBST. To further increase selection pressure, the amount of immobilized target protein was decreased within two subsequent PD iterations while washing stringency was increased during the four panning rounds. Bound phages were cleaved by addition of 30 μg/mL trypsin (Roche Diagnostics, Mannheim, Germany). Phagemids of the selected binding molecules were cloned into pPR-IBAF1b vector (IBA, Goettingen, Germany) to yield expression constructs of the binders with C-terminal *Strep*-tag II. Following transformation of *E. coli* BL21 (DE3) single colonies were picked for screening of candidate binders and cultivated in 96-well scale. Cells were harvested by centrifugation and pellets were suspended in PBST with 250 μg/mL lysozyme (Merck, Darmstadt, Germany) and lysed by three subsequent freeze-thaw-cycles. To select for stable binders the resulting lysates were incubated at 50 °C for two hours leading to heat precipitation of thermodynamically instable variants. After centrifugation screening of soluble EDB-specific binders was performed by ELISA as follows: lysates were incubated with target-coated (67B89) and off-target-coated (6789) 96-well Medisorp-plates (Nunc, Roskilde, Denmark) followed by washing with PBST and PBS. Bound Affilin molecules were detected using an anti-Ubi-Fab-HRP conjugate (AbD Serotec, Puchheim, Germany) using TMB Plus (Kem-En-Tec Diagnostics, Taastrup, Denmark) as substrate. Variants having a target/off-target binding ratio of >2 were defined as binders.

### Affinity maturation, ribosome display and screening of EDB-specific Affilins

Affinity maturation of the precursor variant Af2p, to generate the improved Affilin Af2, was conducted as a sequence of random mutagenesis followed by a selection of binders using ribosome display (RD). The cDNA of Affilin variant Af2p was used as template for error-prone PCR employing the GeneMorph II Random Mutagenesis Kit (Agilent, Santa Clara, CA, USA). The error rate was set to 10-14 mutations per kbp and the resulting theoretical library size was calculated to contain 6·10^10^ independent variants. Linker segments comprising functional elements required for ribosome display were fused via PCR to the 5’ and 3’-ends of the generated library ^41^. Subsequently, four cycles of RD were performed using the PureExpress *in vitro* protein synthesis kit (New England Biolabs, Ipswich, MA, USA) for *in vitro* transcription and translation. Ternary complexes were incubated with the biotinylated target 67B89 in PBSNT (PBS supplemented with 30 mM magnesium acetate and 0.05 % v/v Tween 20) and a 10-fold molar excess of (non-biotinylated) off-target 6789. Target-bound complexes were recovered using M-270 Streptavidin beads (Invitrogen, Carlsbad, CA, USA). Stringent selection of high-affinity binders was achieved by up to 6 washing steps with PBSMT and competitive elution of the immobilized complexes using decreasing concentrations of non-biotinylated target 67B89 in the last two cycles of RD. After each cycle the mRNA was released by addition of EDTA and subsequently reverse-transcribed to obtain the corresponding cDNAs. Prior to the next cycle the cDNA of the pool of binders was re-amplified and supplied with the RD-linker segments by two consecutive PCR reactions. After the fourth cycle of RD the cDNAs of the selected binders were cloned via *Nde*I/*Xho*I restriction sites into an expression vector (pET-Strep-eGFP) providing a genetic fusion of enhanced GFP (eGFP) to the C-terminus of the Affilin molecules for screening. Based on the green fluorescence intensity of single colonies of *E*.*coli* BL21 (DE3) expressing functional Affilin-eGFP fusions without frameshifts, candidates were selected using a K3-XL colony picker (KBiosystems, Basildon, UK) for ELISA binding analysis (as described for the screening after PD). Affilin variants having a target/off-target binding ratio of >2 were defined as binders. For expression without C-terminal eGFP the cDNA of selected Affilins were cloned into the expression vector pPR-IBAF1b (IBA, Goettingen, Germany) providing a C-terminal Strep-tag II.

### Production and purification of Affilin variants

Recombinant protein was expressed in *E*.*coli* BL21 (DE3) in 1-liter scale, followed by purification using a StrepTactin Superflow column (IBA, Goettingen, Germany) according to the manufacturer’s instructions. A second purification step was carried out as size exclusion chromatography on a Superdex 75 pg XK16/600 column, equilibrated in in PBS pH 7.4, using an ÄKTAexpress FPLC system (GE Healthcare, Freiburg, Germany). The tag-free Affilin variants Af1 and Af2 used for binding analysis and crystallization were generated by cloning the respective cDNA into the expression vector pET-20bDoSto, a derivative of pET-20b(+) (Novagen, Darmstadt, Germany) providing a C-terminal hexahistidine tag and an additional stop codon. Plasmids were subsequently transferred into electro-competent *E. coli* BL21 (DE3) cells. Expression was carried out in 1 L scale. After cell harvest and cell disruption, proteins were purified from lysates via a HiTrap Q Sepharose FF anion exchange column and subsequent HiTrap Phenyl HP hydrophobic interaction chromatography. Purified Affilin variants were then dialyzed against PBS pH 7.4.

### Production of Interleukin-2 fusions

EDB-specific Affilin variant Af2 and the non-binding control diUb-WAA were genetically fused to the N-terminus of a synthetic gene of human Interleukin-2 (UniProtKB ID P60568, residues 21-153) separated by a 15 amino acid (Ser_4_-Gly)_3_ linker. The constructs were cloned into the vector pET-28aS, a derivative of pET-28a (Novagen, Darmstadt, Germany), via *Bsa*I*-*HF restriction sites, providing a tag-free expression. Resulting plasmids were transferred into electro-competent *E. coli* BL21 (DE3) cells and protein expression carried out for 4 hours in 1 L scale at 37°C. Insoluble protein expression required *in vitro* refolding of Af2-IL-2. Based on established protocols ^42 43^, harvested cells were disrupted and inclusion bodies were isolated and solubilized in 6 M guanidine hydrochloride, 100 mM Tris pH 8.5, 1 mM EDTA, 100 mM DTT. Renaturation was carried out by rapid dilution pulses of the solubilized protein (final concentration 100 μg/mL) into 1 L buffer containing 50 mM Tris pH 9.0, 3 M urea, 2.5 mM GSH, 0.25 mM GSSG) at 4°C under gentle stirring for 16 hours. Subsequent purification of the refolded protein was achieved by addition of (NH_4_)_2_SO_4_ (1 M final concentration) followed by filtration, purification via hydrophobic interaction chromatography (HiTrap Phenyl HP column) and size exclusion chromatography (XK26/600 Superdex 75 prep grade, equilibrated in PBS pH 7.4).

### Binding analysis by Surface Plasmon Resonance (SPR)

Surface plasmon resonance measurements on a Biacore 3000 (GE Healthcare, Freiburg, Germany) were used to analyze binding of purified Affilin variants to the target 67B89. Purified biotinylated target was immobilized on a streptavidin chip (GE Healthcare) according to the manufacturer’s instructions. The off-target 6789 was immobilized to the reference channel of the chip. Different concentrations of the Affilin variant (0 - 100 nM) were analyzed for binding to 67B89 using PBS pH 7.4 containing 0.005 % Tween 20 as running buffer at a flow rate of 30 μl/min. Traces were corrected by subtraction of the reference signal and the trace of buffer injection. K_D_, k_on_ and k_off_ values were calculated by fitting the traces using a global kinetic fitting (1:1 Langmuir model, BIAevaluation software).

### Binding analysis by ELISA

Determination of binding affinity by ELISA was carried out in 96-well medium binding plates (Microlon 200, Greiner Bio-One, Kremsmuenster, Austria). Plate coating with 5 μg/ml target 67B89 (or the target variants 67B, 7B8, B89, respectively) and off-target 6789 was performed by overnight incubation at 4 °C. The wells were washed three times with PBST and blocked with 3 % BSA solution for 2h at 20°C. Binding to the target-coated plate was performed in concentration-dependent manner by incubation with the binder at concentrations up to 100 nM followed by washing the plates three times with PBS. Bound Affilin molecules were detected using an anti-Ubi-Fab-HRP conjugate (AbD Serotec, Puchheim, Germany) using TMB Plus (Kem-En-Tec Diagnostics, Taastrup, Denmark) as substrate. POD activity was measured spectrophotometrically at a wavelength of 450 nm in a microplate reader (Sunrise, Tecan, Maennedorf, Switzerland). Binding to the target 67B89 was determined in triplicates and by single measurement for the off-target 6789, respectively.

### Analysis of thermal stability

Thermal unfolding transitions of proteins were measured by means of differential scanning fluorimetry (DSF), performed at a protein concentration of 0.1 mg/ml protein in PBS pH 7.4 using a 10-fold dilution of SYPRO Orange (Invitrogen, Carlsbad CA, USA) in a real-time PCR device (Light Cycler 480, Roche Diagnostics, Mannheim, Germany). Fluorescence was recorded at 465 nm excitation and 580 nm emission wavelengths, respectively, over a temperature range of 20-90°C with 1 K/min heating rate. For evaluation, fluorescence intensity was plotted against the temperature and the inflection point (T_m_) derived from the maximum of the first derivative of the plot as the midpoint of thermal unfolding.

### Cell binding analysis by immunofluorescence

The Affilin variants Af2s and Af2-IL2 were analyzed for binding to EDB expressing Wi-38 cells (human fetal lung fibroblasts, ATCC CCL-75) using NHDF cells (neonatal human dermal fibroblasts, Promocell, Heidelberg, Germany), lacking EDB expression, as negative control. Cells (30,000 per well) were disseminated and cultivated for 96 hours in 4-well chamber slides in 90% EMEM medium supplemented with 10% FBS (Wi-38 cells) and 98% Fibroblast Growth Medium/2% supplement mix (Promocell, NHDF cells), respectively. After washing three times with PBS, fixation with methanol, washing and blocking (5% horse serum in PBS) cells were incubated with 50 nM Af2s and Strep-tagged diUb-WAA (as non-binding control), respectively, for 1 h at 37 °C. For detection of bound Af2s, a rabbit anti-Strep-tag IgG antibody (Genscript, Piscataway, NJ, USA) and a goat anti-rabbit IgG Alexa 488 conjugate as secondary antibody were used. Detection of bound Af2-IL was performed using a rat anti-human IL-2 mAb Alexa Fluor488 conjugate (Invitrogen, Carlsbad CA, USA). Cell nuclei were counterstained with DAPI (Sigma Aldrich, Steinheim, Germany) and embedded in the polyvinyl alcohol Mowiol. Visualization was conducted using an Axio Scope AF1 fluorescence microscope (Zeiss, Jena, Germany) employing EX BP 470/40, BS FT 495 and EM BP 525/50 filters. For DAPI visualization filters EXG365, BS FT 395 and EM BP 445/50 were used.

### Activity assay of Interleukin-2 fusions

The IL-2 activity assayed using murine cytotoxic T-cells (CTLL-2, ATCC TIB-214) dependent on IL-2 for growth ^44^. Cultivation was performed in full medium (78% RPMI1640, 10 % FBS, 10 % T-STIM, 2 mM L-glutamine, 1 mM sodium pyruvate). Harvested cells were washed twice with medium without T-STIM. Subsequently, 40,000 cells were seeded per cavity of a 96-well plate and incubated for 20 h with serial dilutions (1000 – 0.076 pM) of the fusion protein Af2-IL-2 and the reference recombinant human IL-2 (PeproTech, Rocky Hill NJ, USA), respectively. Viable cells were detected using WST-1 reagent (Roche Diagnostics, Mannheim, Germany) by absorption measurement at a wavelength of 450 nm. EC_50_ values from triplicate measurements were calculated using the program PLA 2.0 (Stegmann Systems, Rodgau, Germany).

### Protein crystallization and structure determination

Complex formation of tag-free Affilin variants Af1 and Af2, respectively, with the truncated target 7B8 was accomplished by incubation of equimolar concentrations of the binding partners at 20°C for 1 h. The complex was isolated from unbound species by size exclusion chromatography (Superdex 75 26/600) in 10 mM HEPES, 100 mM NaCl pH 7.3 and concentrated to 22 mg/mL using Amicon Ultra-4 centrifugal filters (Millipore, Billerica MA, USA).

Unbound Af1 was crystallized during attempts to obtain complex crystals of Af1 with the target 7B8. Small bipyramidal crystals were obtained within 2 weeks from hanging drop vapour diffusion crystallization setups of the Af1:7B8 complex at 15°C in 100 mM Imidazol/MES, pH 6.0, 60 mM calcium/magnesium chloride, 30% PEG 8000/ethylene glycol (w/v) including 10 mM copper(II)chloride dehydrate. Initial crystals were used for macro-seeding to grow larger crystals of the same morphology suitable for X-ray diffraction analysis. Data collection from a single frozen crystal was carried out at beamline 14.2 at the BESSY II electron storage ring (Helmholtz-Zentrum für Materialien und Energie, Berlin; Germany). A 2.2Å dataset was collected at a wavelength of 0.9184 Å using a CCD detector (MX-225, Rayonics, USA). Diffraction data were processed with the XDS software package ^45^. The structure was phased by single wavelength anomalous dispersion (SAD) resulting in the localization of 4 heavy atoms (copper) in space group P4_1_ employing the SHELX software suite ^46^. Subsequent cycles of heavy atom refinement and density modification, carried out with the software AUTOSHARP ^47^, yielded an initial electron density which allowed automated tracing of the polypeptide backbone using ARP/wARP from the CCP4 suite ^48, 49^. The model was completed by manual building using the program Coot ^50^ and refined with the PHENIX software suite ^51^. The structure revealed one Af1 molecule in the asymmetric unit. The electron density was well resolved for most residues of Ub-N and Ub-C, except for the side chains of several evolved residues in the solvent-exposed α2β5 loops (N62, P63, K64, L65, W142, Q143) and the linker region not being visible (A76, G77, I78, G79).

Crystals of the Af2:7B8 complex grew within 1-2 weeks in sitting-drop vapor diffusion plates at 25°C in 500 mM lithium sulfate, 15% PEG 8000 (w/v). Diffraction data were collected from a single frozen crystal at the BESSY II beamline BL 14.1 using a hybrid pixel detector (Pilatus 6M, Dectris, Switzerland) and processed with the XDS software package. Phases were determined by Molecular Replacement employing the program PHASER ^52^ using the three (separated) fibronectin domains of 7B8 (PDB entry: 4GH7 ^53^) and the two (separated) ubiquitin domains of Af1 as individual search models. Three non-crystallographic symmetry-related complexes were located in the asymmetric unit, each consisting of one Af2 and one 7B8 molecule. The structure was completed using the program Coot ^50^ and refined with the PHENIX software suite ^51^. Structure validation was carried out using MOLPROBITY ^54^, molecular figures were created with the software PyMOL (Schrödinger LLC, New York, USA). Data collection and refinement statistics are given in supplementary table S1.

### Analytical HPLC size exclusion chromatography

Analytical size exclusion HPLC of purified complexes of Af1:7B8 and Af2:7B8 were carried out using a Superdex 200 5/150 GL column (GE Healthcare, Freiburg, Germany) equilibrated in PBS, using a Summit HPLC system (Dionex, Idstein, Germany). The apparent molecular masses were derived from a calibration of the column by linear regression of the retention times of an HPLC gel filtration standard (Bio-Rad, Feldkirchen, Germany) and were compared to theoretical masses of the complexes (47.3 and 47.2 kDa, respectively), calculated from the individual molecular masses (Af1: 17.5 kDa, Af2: 17.4 kDa, 7B8: 29.8 kDa).

## Footnotes

## Author contributions

A.K., E.F., U.H. and C.P. designed experimental work, A.K. performed biochemical experiments and crystallization, A.R., F.S. and M.G.-B. carried out biochemical and cell biological experiments, C.P. solved structures and performed structural analysis, A.K., E.F. and C.P. wrote the manuscript.

## Conflict of interest statement

A.K., E.F., U.H., F.S., M.G.-B. are employees of Navigo Proteins GmbH. Affilin, Navigo, Navigo Proteins are registered trademarks of Navigo Proteins GmbH.

### Data deposition

The atomic coordinates and structure factors of unbound Af1 and the Af2:7B8 complex have been deposited in the Protein Data Bank, www.pdb.org (PDB ID codes 8PF0 and 8PEQ, respectively).

## Acknowledgements

We thank Milton T. Stubbs for helpful discussions, the Helmholtz-Zentrum Berlin für Materialien und Energie for the allocation of synchrotron radiation beam time and the staff of the MX beamlines at the BESSYII storage ring for excellent support.

## Display items

**Extended data figure 2.**
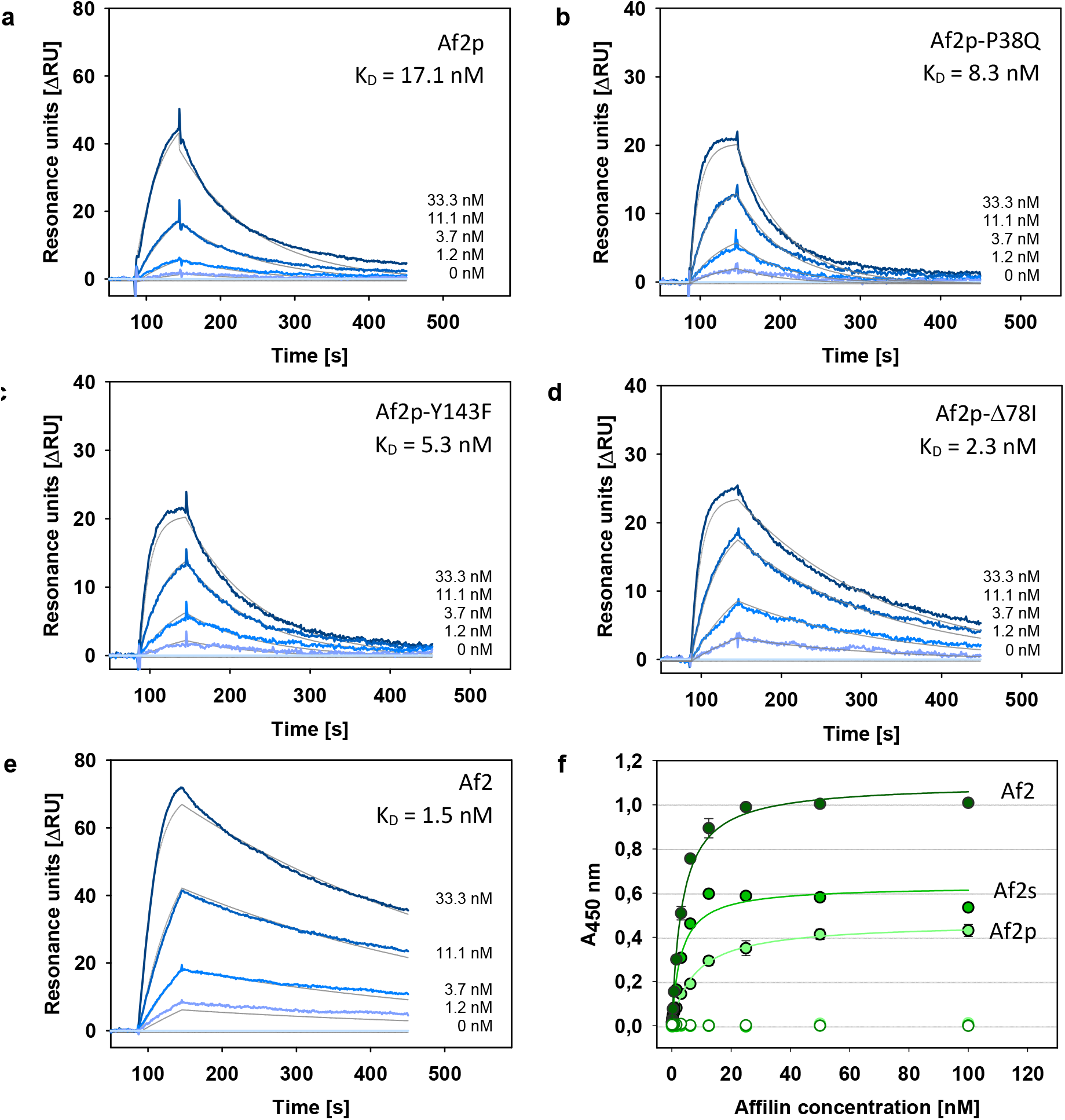
Binding analysis of Affilin variants generated during the directed evolution process with the target 67B89 by SPR or ELISA (complete binding parameters are given in table 1). (**a**) Precursor variant Af2p generated by phage display. (**b**), (**c**), (**d**) Single amino acid exchange variants of Af2p, individually introducing one of the three mutations acquired during affinity maturation of Af2p. (**e**) Strep-tag-free variant Af2, crystallized in complex with 7B8. (**f**) Binding of Affilin variants to the target 67B89 (filled circles) and lack of binding to the off-target 6789 (open circles) analyzed by ELISA.

**Extended data figure 2.**
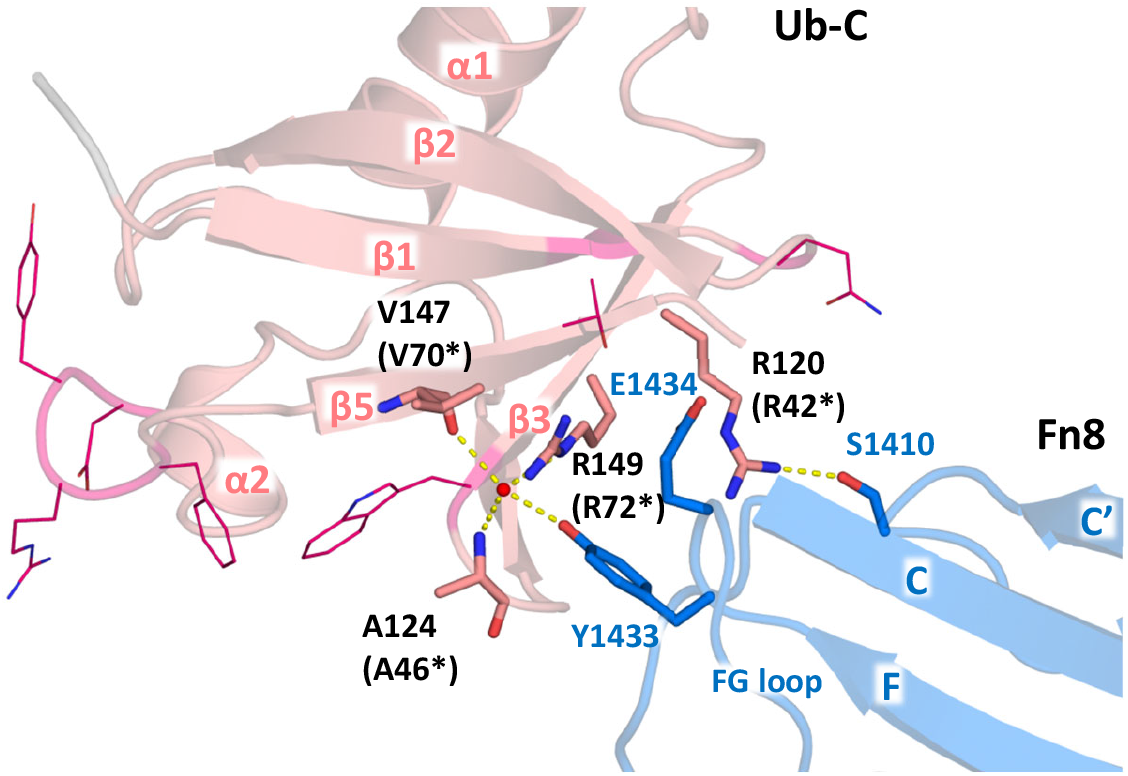
Molecular interactions contributing to FN8 binding by Af2 Ub-C exhibiting a -4 register shift in β5. Residues coloring of C-atoms in magenta/light red (Ub-C), blue (Fn8), white (Ub-wt), N-atoms in blue and O-atoms in red. Detailed view of key interaction between Ub-C and EDB (interface III), residues involved in binding shown as sticks, evolved amino acids of Ub-C depicted as lines, hydrogen bonds shown as yellow dashed lines.

**Extended data figure 3.**
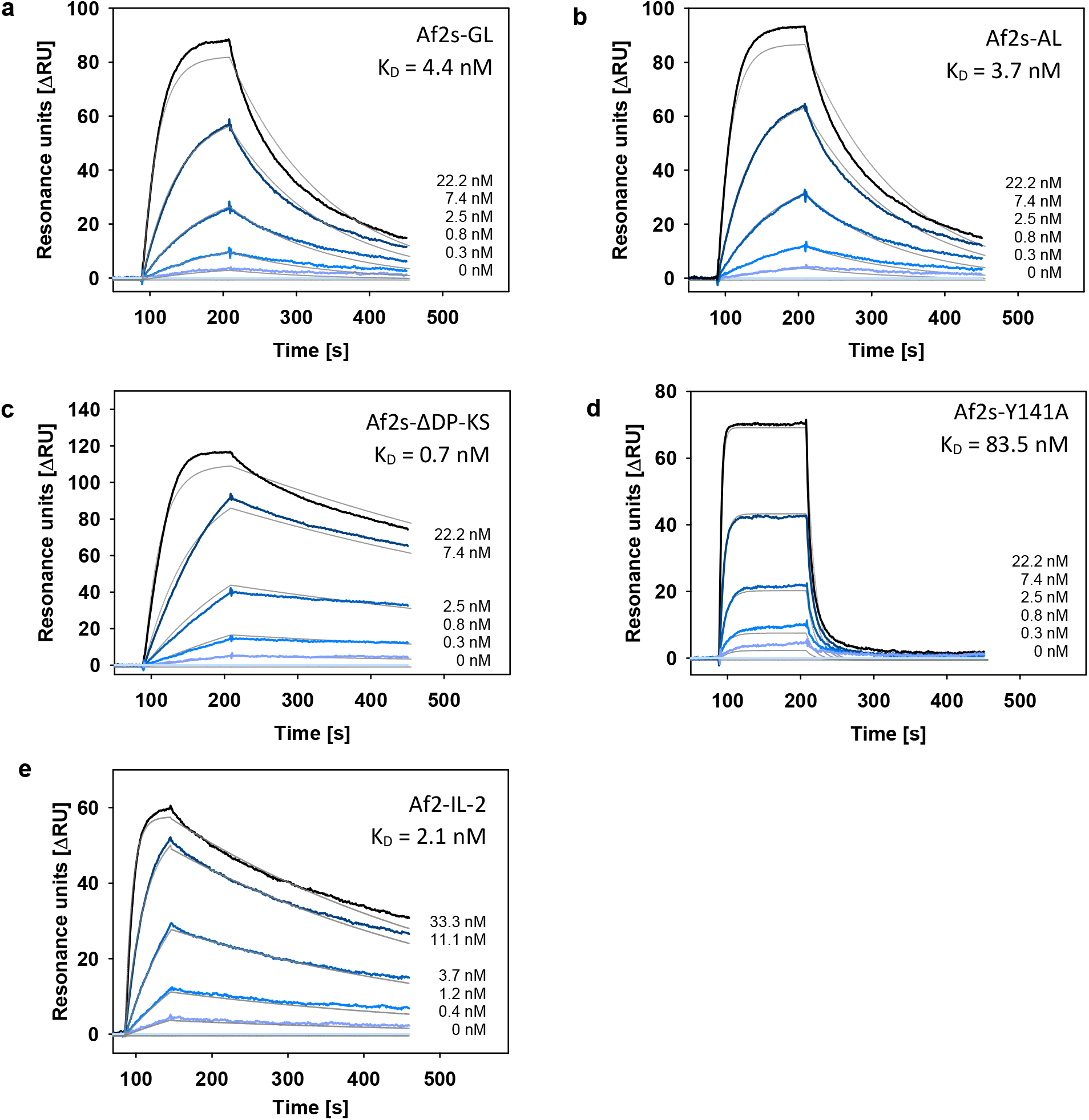
Influence of selected modifications on the target binding properties of Af2s analysed by SPR at different Affilin concentrations (complete binding parameters are given in table 1). (**a**) Variant Af2s-GL carrying the mutations R74*G and α75*L in Ub-C. (**b**) Variant Af2s-AL carrying the mutations R74*A and α75*L in Ub-C. (**c**) Variant Af2s-ΔDP-KS carrying the deletions ΔD62 and ΔP63 and the mutations L65K and L67S in Ub-N. (**d**) Variant Af2s-Y141A (Y64*A). (**e**) Variant Af2-IL-2, genetic fusion of Af2s to human IL-2.

**Extended data figure 4.**
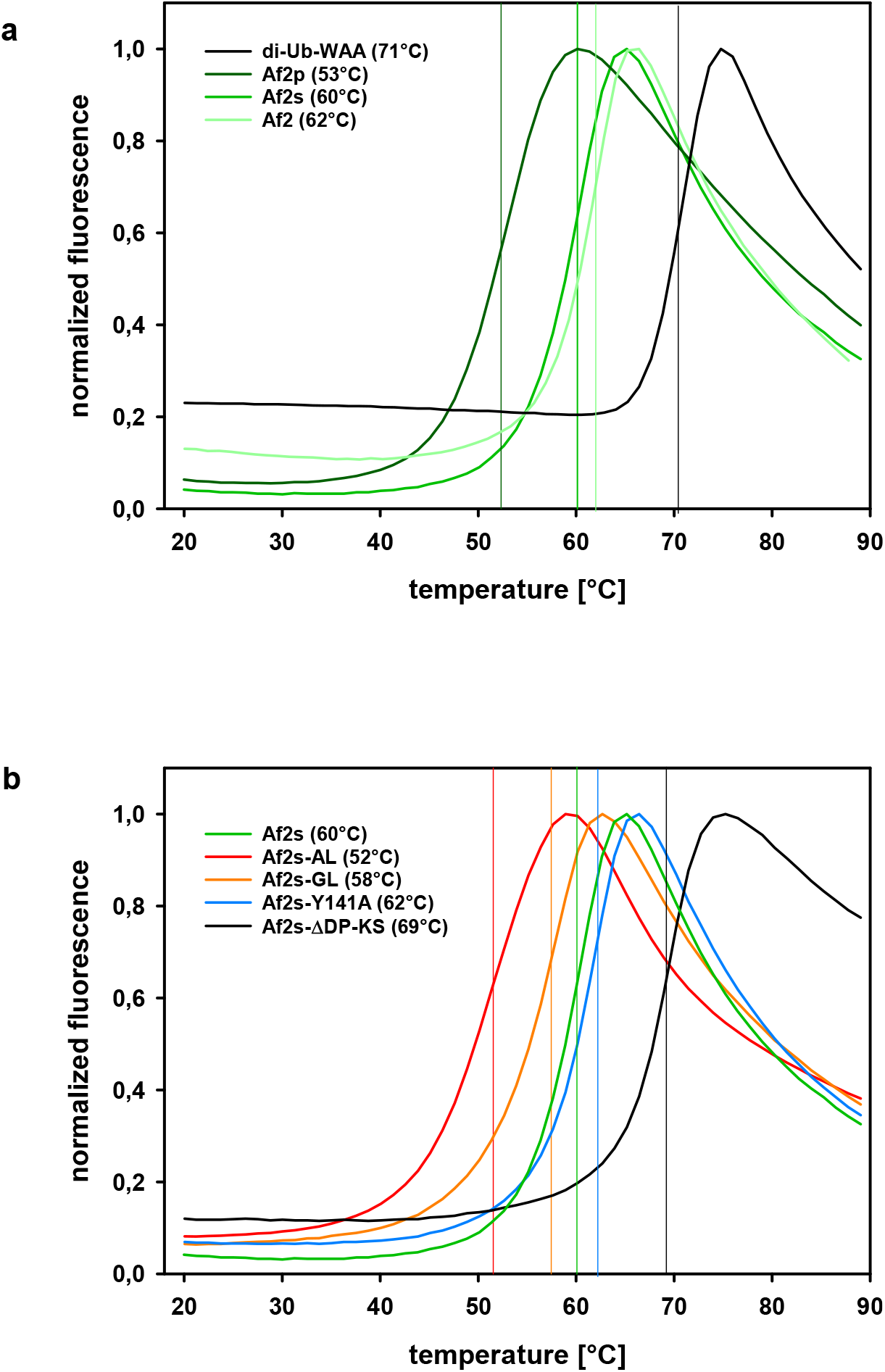
Analysis of thermal stabilities of Affilin variants by DSF (**a**) Variants Af2p, Af2s and Af2 in comparison with a non-evolved diubiquitin variant (diUB-WAA). (**b**) Variant Af2s-GL carrying the mutations R74*G and α75*L in Ub-C, variant Af2s-AL carrying the mutations R74*A and α75*L in Ub-C, variant Af2s-ΔDP-KS carrying the deletions ΔD62 and ΔP63 and the mutations L65K and L67S in Ub-N and variant Af2s-Y141A (Y64*A). Vertical lines mark the midpoints (T_m_) of thermal unfolding.

**Extended data figure 5.**
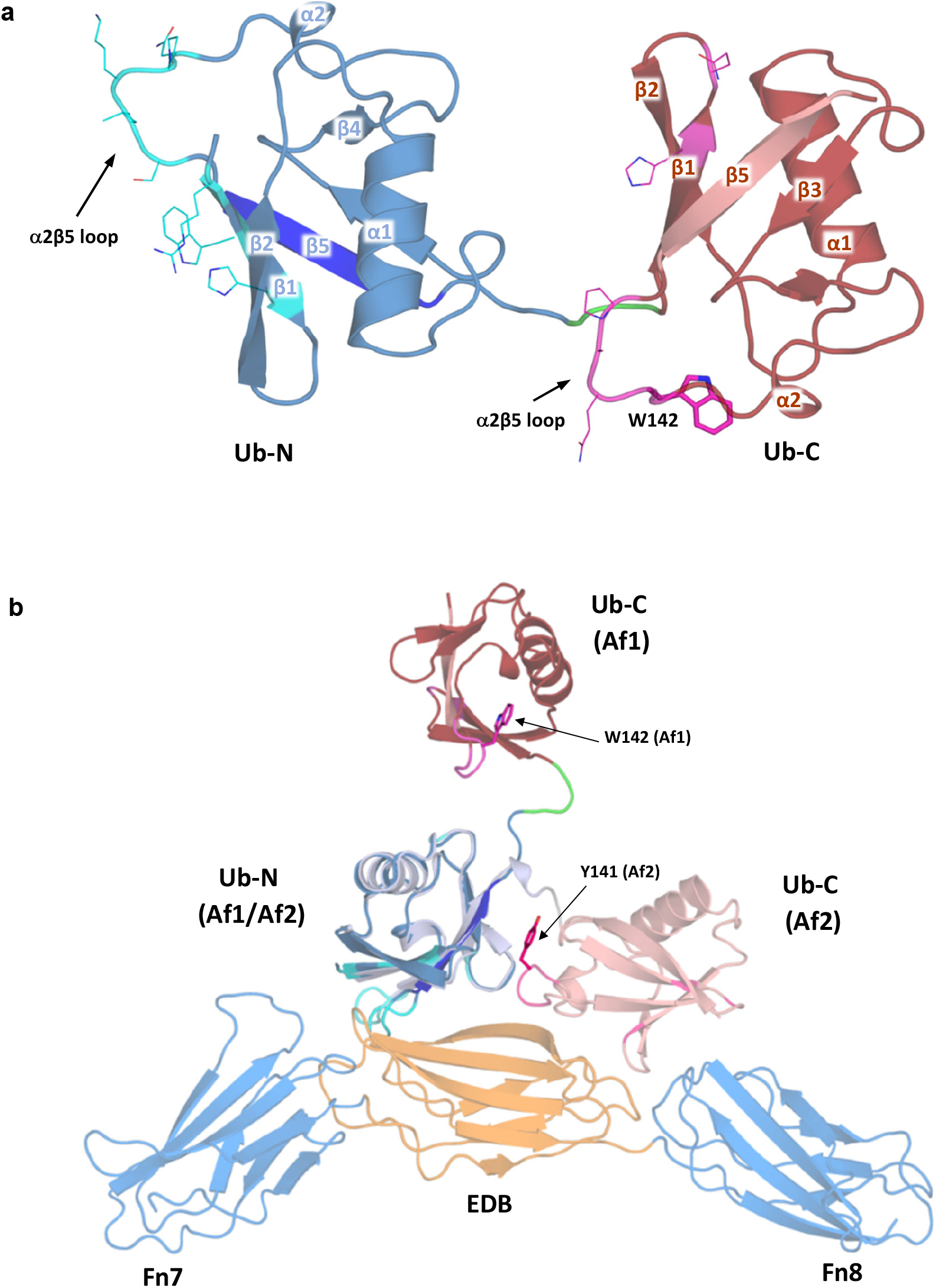
Crystal structure of unbound Affilin Af1 **(a)** Overall structure of Af1 highlighting the secondary structure elements with β5-strands of Ub-N (dark blue) and Ub-C (light pink), evolved residues depicted as lines, colored by atom type (C-atoms of Ub-N cyan, C-atoms of Ub-C magenta, N atoms blue, O atoms red). Linker residues colored in green. W142 (corresponding to Y141 in Af2) shown as sticks. (**b**) Structural superposition of Af1 and the Af2:7B8 complex based on the alignment of the Ub-N domains. Regions of evolved residues colored in cyan (Ub-N) and magenta (Ub-C). Y141 (Af2) and W142 (Af1) shown as sticks.

