## Supplemental Information for "Ubiquitin-derived artificial binding proteins targeting oncofetal fibronectin reveal scaffold plasticity by β-strand slippage"

to

Katzschmann *et al.*

#### Supplementary discussion

##### Comparison of the directed evolution strategies for Af1 and Af2

The high binding affinities and specificities of Af1 and Af2 towards oncofetal fibronectin fragments, as well as their thermal stability, result from similar directed evolution approaches relying on the same diUb scaffold starting point (variant diUb-WAA). Af1 [1] and Af2 (this work) were evolved by combinatorial library design randomizing selected amino acid positions, two stages of stringent selection using phage and ribosome display techniques followed by extensive candidate screening. Nevertheless, the strategies differed in several apects: both Affilin molecules originate from independently prepared cDNA libraries, randomizing the same set of 14 residues from β-strands (β1, β5) and loop regions (β1β2, $\alpha$2β5) of Ub-wt (with two additionally permutated amino acid positions in strand β1 of Af1 Ub-N) (figure 1e, extended data figure 5a, supplementary figure S1). In case of Af2, a selection for thermal stability (>50°C) was introduced, following the first selection stage (phage display, PD), yielding the precursor variant Af2p. Another important difference was the strategy used for affinity maturation: for Af2p random mutagenesis by error-prone PCR was employed followed by a selection step, involving a C-terminal genetic fusion of eGFP to the candidate cDNA, allowing fluorescent screening of properly folded candidates, thereby excluding false-positive binders generated by frame-shift mutations. In contrast, for affinity maturation of Af1 the previously selected residues of the C-terminal domain (Ub-C) were re-randomized, while the sequence of the Ub-N was preserved. Similar for Af1 and Af2, Affilin candidates were enriched in a second selection step by ribosome display (RD), increasing stringency with every iteration, resulting in higher binding affinities and specificities as well as enhanced thermal stabilities of the Affilin proteins. Albeit Af1 and Af2 were both raised against the same target 67B89, the majority of the evolved amino acid substitutions differ between Af1 and Af2 (exceptions are H6, P63, L65 and Q86, supplementary figure S1).

##### Contribution of individual mutations evolved during affinity maturation of Af2

Maturation of the parental variant Af2p, yielding Af2s (the Strep-tagged version of Af2), resulted in a further increase of affinity towards 67B89, as shown by the low nanomolar $K_D$ values from ELISA and SPR analysis (table 1, extended data figure 1). Interestingly, the increase of affinity during maturation seems to arise mostly from a 5-10-fold lower off-rate $k_{off}$ for Af2s/Af2 compared to Af2p, while the on-rate $k_{on}$ is only moderately affected. The striking recovery of thermal stability by the increase of +9K in $T_m$ (table 1, extended data figure 4a) during maturation of Af2p to Af2 is a consequence of the three sequence alterations (two mutations P38Q, Y143F and one deletion of I78). To further address the contribution of the individual events, three Af2p variants (Af2p-P38Q, Af2p-Y143F and Af2p-$\Delta$78I) were generated, purified and subjected to binding analysis by SPR (table 1, extended data figure 1). Measurements indicated a cumulative effect: each individual sequence change occurring in the maturation process seems to contribute to higher affinity (lower $K_D$), by increasing the rate of association ($k_{on}$), while shortening the linker between the two Ub domains by deletion of Ile78 leads also to a decrease of the dissociation rate ($k_{off}$). Apparently, from the three mutations $\Delta$78I has the largest contribution to the gain of affinity. Presumably, reducing the linker length between Ub-N and Ub-C places both domains in optimal distance for target binding and limits conformational freedom of the domains relative to each other. With a restricted arrangement, both Ub domains can then act as a single, almost rigid binding module, rather than to two flexibly tethered binding domains. The benefit would be reduced entropic costs of target binding and could lead to the observed gain of affinity of Af2 variants, larger than a sole avidity effect of two tethered Ub-domains would produce [3].

**Description of the asymmetric unit of the Af2:7B8 crystal structure**

The crystal structure of the Af2:7B8 complex elucidated from monoclinic crystals, revealed three complexes in the asymmetric unit, related by a $3_1$ non-crystallographic symmetry-axis (supplementary figure S3). Each copy represents a heterodimer formed by one Af2 chain and one 7B8 chain, respectively, confirming the 1:1 stoichiometry suggested by HPLC-SEC analysis (supplementary figure S2). Both Ub-N and Ub-C display the canonical β-grasp fold of Ubiqutin-like domains, composed of a twisted β-sheet formed by five antiparallel β-strands (β1 – β5) embedding an α-helix (α1) and a short $3_{10}$-Helix (α2) (figure 1e). The domain orientations of Ub-N and Ub-C relate to each other by a rotation of about 60°. From the target, Fn7, EDB and Fn8 contain two β-sheets forming the typical β-sandwich structure of Fn type III domains [4], [5]. Three antiparallel β-strands (A, B, E) constitute the first β-sheet and pack against the second sheet, consisting of four antiparallel strands (C, C', F, G). The two tethered Ub-like domains of Af2 are positioned in an antiparallel orientation to the Fn domains of 7B8 and thereby fix the angle of the hinge region between EDB and Fn8 in the target and also the flexible linker between Ub-N and Ub-C in the Affilin.
Superposition of the three Af2:7B8 complexes of the asymmetric unit (by structural alignment of the EDB domains) resulted in higher root-mean-square deviation (RMSD) for the Fn7 domains (average $C_\alpha$ RMSD 7.0 Å) than for the EDB and Fn8 domains (average $C_\alpha$ RMSD 0.8 Å and 1.8Å, respectively) (supplementary figure S3c). As Fn7 is not directly involved in binding, its position is less restricted, with the Fn7-EDB linker region acting as a hinge. Only the target domains EDB and Fn8 interact with Af2 in very similar fashion within the three complexes, judged by the positions of the three superimposed Affilin molecules relative to their target ($C_\alpha$ RMSD 0.9 Å). Notably, the position of one Af2 chain deviates more (chain M, average $C_\alpha$ RMSD 2.6 Å) from the other two (chains L and J) in the asymmetric unit. This difference originates mainly from packing effects between the complexes within the asymmetric unit, resulting e.g. in altered conformations of the β1β2 loop of Ub-N and the linker region between Ub-N and Ub-C of chain M (supplementary figure S3c). The structural description of the Af2:7B8 complex refers to the complex formed by chain A (7B8) and L (Af2).

**Detailed description of the Af2 target-binding interface**

In the complex structure of Af2:7B8, Ub-C contributes the major part (interface II, 485 Å and III, 326 $Å^2$) to the target binding interface compared to Ub-N (interface I: 368 $Å^2$). From 7B8, only the EDB and the Fn8 domain contribute to the binding interface, predominantly the EDB (interfaces I + II) , the Fn8 domain (interface III) to a lesser extent, while the Fn7 domain appears not to be directly involved in the interaction (figure 1d).
From interface I (figure 2a), residues F4 and evolved H6 from β1 of Ub-N are in hydrophobic interaction to EDB residues I1296 and F1315 (by π-stacking), respectively. The sidechains of Ub-N K66 (evolved, α2β5 loop) and H68 (non-evolved, β5) interact via hydrogen bonds to backbone atoms of EDB I1295 and I1296. The carboxyl moiety of EDB D1317 sidechain forms an intermolecular hydrogen bond to the backbone amide of Ub-N K66, while also electrostatic interaction between the sidechains of both residues are conceivable.
Key interactions of interface II (figure 2b) comprise a salt bridge between the sidechains of EDB D1314 and Ub-C R142 (evolved) and hydrophobic interactions of EDB F1312 to evolved F143. Both Ub-C residues are located in the α2β5 loop. Also the guanidine moiety of Ub-C R149 forms (partially water-mediated) hydrogens bonds and a salt bridge to the sidechain of EDB E1329. Several further contacts contribute to the intermolecular bond network of sub-site II: e.g. van-der-Waals interactions between Ub-C W123 and the EF loop region around G1327 of EDB.
Interface III (extended data figure 2) employs only non-evolved residues: R120 (β3 strand), A124 (β3β4 loop) and the residues V147 and R149 (β5 strand) interact via (partially water-mediated) hydrogen bonds to Y1433 and S410 of Fn8, respectively. Additionally R149 forms a salt bridge to E1434 (FG loop) of Fn8.

The target 7B8 exhibits an overall acidic character (calculated pI=4.1) and reveals a significant negative electrostatic potential at the surface (supplementary figure S4), while the Affilin Af2 (calculated pI=5.6) displays a relatively positive electrostatic potential at the binding interface to 7B8. This charge complementarity of the Af2:7B8 interface suggests a contribution of ionic interactions, adding to the framework of intermolecular hydrogen bonds, hydrophobic and van-der-Waals interactions, involved in target binding.

**Structural basis of the -2 register-shift in the Ub-N domain of Af2**

In addition, several intramolecular interactions within Af2 as well as intermolecular interactions to the bound target contribute to stabilisation of the register-shifts observed in Ub-N and Ub-C that include evolved as well as conserved residues, partially recruited into new environments by the register shifts. Intramolecular interactions include hydrophobic clusters e.g. formed by W45, I61, L67 and L69. While this cluster can also be formed in the non-shifted conformation (with L65 and L67 in place of L67 and L69, respectively), a second hydrophobic cluster, formed by I44 and V70, the $\pi$-stacking interactions of F4 and H68 and an intramolecular hydrogen bond between H68 and the evolved residue H6 can only occur in the -2-shifted state(supplementary figure S7a). The non-shifted conformation would also be destabilized by the evolved charged residue D8 of Af2, precluding the hydrophobic interactions of L8 to V70, found in the structure of of Ub-wt.
The bound target contributes to further stabilization of the register-shifted conformation of Ub-N, e.g. by the electrostatic interactions (D1317 of 7B8 to evolved K66 of Af2 Ub-N) and hydrogen bonds (between the sidechain of H68 and the backbone carbonyl oxygen of I1296). These interactions can occur only in presence of the -2 register shift, bringing the respective residues into spatial proximity (figure 2a).
The $\alpha$2β5 loop extension, induced by the -2 register shift, could potentially impose sterical problems to this region of Ub-N with – as observed - detrimental effects on Affilin stability and binding affinity. To assess this further, the variant Af2s-ΔDP-KS was generated. It comprises a deletion of two residues (D62 and P63) from the $\alpha$2β5 loop, and two additional mutations to restore original Ub-wt residues: loop residue L65 is reverted to lysine and L67 back to serine (supplementary figure S1). Thus, K65 and S67 of Af2s-ΔDP-KS should occupy the original positions of K63 and S65 in Ub-wt, avoiding hydrophobic residues in the $\alpha$2β5 loop and at the S65-binding pocket, while the -2 register shift of β5 should be preserved. Evolved K66 (T66 in Ub-wt) was not substituted, as it is involved in EDB binding (figure 2a). Variant Af2s-ΔDP-KS revealed a remarkable increase of thermal stabiliy ($\Delta T_m$=+9 K) compared to Af2s, and binding analysis by SPR revealed an about 5-fold higher affinity to the target 67B89 (extended data figures 3c and 4b, table 1). The observed gain of affinity resulted predominantly from an increased association rate ($k_{on}$). The data suggest that the -2 register-shifted state of Af2 Ub-N can be further stabilized by shrinking the extended $\alpha$2β5 loop size again and by “reversal” of residue dislocation at β5. Unfortunately, efforts to crystallize the variant Af2s-ΔDP-KS remained unsuccessful.

**Structural basis of the -4 register-shift in the Ub-C domain of Af2**

The four evolved residues in the extended $\alpha$2β5 loop of Af2 Ub-C stabilize the -4 shift by several intra- and intermolecular interactions (supplementary figure S7b). The sidechain of F66* is buried in the hydrophobic cluster of W45*, L69*, V70* (asterisks designating residue numbering corresponding to Ub-N). An intra-loop salt bridge (and two hydrogen bonds) between R65* and D63* and the locked sidechain of Y64* (Y141), being sandwiched between the Ub-N and Ub-C domains, restrict the conformation of the $\alpha$2β5 loop, further stabilizing the -4 shifted conformation (figure 1e). Intermolecular interactions to the bound target 7B8 also promote the -4 register shift: the electrostatic interactions of R65* and R72* to EDB residues D1314 and E1329, respectively, as well as the hydrophobic interaction between F66* and F1312 of EDB all involve Ub-C residues in register-shifted positions (figure 2e). As in Af2 Ub-N, the two hydrophobic grooves of Ub-C are occupied by leucines, but now by L71* and L73* (supplementary figure 6d). The β5 strand terminates at R72* (located at the position of H68 in Ub-wt), leaving a potential third binding site, harboring L71 in Ub-wt and L73 in Ub-N, unoccupied. To assess the relevance of a third leucine binding groove, two Af2s variants were generated: Af2s-GL and Af2s-AL. In both variants A75* is replaced by L75*, which could re-occupy the potential third site with a leucine (supplementary figure S1). Furthermore, in both variants R74* was replaced with either G74* or A74*, respectively, to reduce any potential sterical restrictions from the bulky side chain of Arg in its shifted position. Thermal stabilities of the Af2 variants and binding to the target 67B89 were analysed (table 1, extended data figures 3 and 4b). The amino acid changes showed only moderate effects: while the thermal stabilities were decreased (Af2s-GL by 2 K and Af2s-AL by 8 K), the target binding affinities remained almost unchanged, in spite of higher rates of association ($k_{on}$) and dissociation ($k_{off}$). This defines only the L67/L69-binding grooves of ubiquitin to be relevant, allowing register shifts by 2 or 4 residues, where in all three states of β5 (non-shifted, -2-shifted, -4-shifted), the both leucine-binding grooves are occupied by leucines residues (supplementary figure 6c-d).

Interestingly, the extension of the α2β5 loop of Af2 Ub-C by four residues from β5 coincides with the deletion of a residue (ΔQ140) from the same loop during the evolution of Af2 by phage display, resulting in an extension of α2β5 by only three residues (supplementary figure S1). Presumably, the deletion counteracted the -4 register shift, by partially compensating the space requirements of the four additional amino acids to be accommodated in the loop. Similar findings were previously reported for a mono-ubiquitin variant (Ubv-10F), evolved to bind tumor necrosis factor TNFα [6]. Two amino acids (D58, Y59) at the N-terminal boundary of the α2β5 loop region were found to be deleted during evolutionary selection of Ubv-10F. Re-introduction of both residues at their original location abolished binding of the ubiquitin variant to the target TNFα. However, without a three-dimensional structure of Ubv-10F it can only be speculated, that a β5 register shift is also present in Ubv-10F, defining a prerequisite for target binding.

**Structural basis of the -2 register-shifts in Ub-N and Ub-C of Af1**

With a Cα RMSD of 0.2Å, the overall domain structures of Af1 Ub-N und Ub-C are very similar to each other. Also the α2β5 loops of both domains adopt a nearly identical backbone conformation, although they comprise different sets of evolved residues (supplementary figure S8). Interactions stabilising the -2 shift in Ub-N and Ub-C of Af1 strongly resemble also the interactions supporting the -2 register shift in Af2 Ub-N. The evolved residues L65 (Ub-N) and A65* (Ub-C) are displaced from their original position in β5, now inhabited by L67, contributing to the hydrophobic interactions clustered around W45 (involving also I61 and L69, identical residues in Ub-C not mentioned). V70, now -2-shifted to the former position of H68 in β5, forms an additional hydrophobic patch with I44 on the opposite face of the β-sheet, while H68 could form π-stacking interactions to an aromatic residue (W4, F4*) in the neighbouring β1 strand. Interestingly, for both Ub domains of Af1 the register shifts appear to be stabilized by chelation of copper ions from the crystallisation solution, which coordinate the imidazole side chains of H6 (β1) and H68 (β5), while in Ub-N of Af2 the same residues share a hydrogen bond instead (supplementary figure S7a).

**Structural comparison of Af2 Ub-N and Ub-TVLN**

The structure of Ub-TVLN bound to PINK1 [7] is strikingly similar to Ub-N of Af2 bound to 7B8. Both structures display a -2 register shift and with an overall Cα RMSD value of 0.8 Å even the backbone conformation of the α2β5 loops are nearly superimposable, although their amino acid composition differs substantially. Interestingly, identical subsets of amino acid positions (e.g. residues 65, 66 and 68 in the α2β5 loop) are involved in interactions of Af1 Ub-N and Ub-TVLN to very different binding partners, oncofetal fibronectin and the Ub kinase PINK1, respectively (supplementary figures S9d-e and h). This is noteworthy, as the Affilin library design was conducted before elucidation of the PINK1:Ub-TVLN structure, hence unbiased from this structural knowledge.

**Structural comparison of Af2 Ub-C and Ubv-G08**

Superimposition of Af2 Ub-C and 53BP1-bound Ubv-G08 [8] (overall Cα RMSD 1.4Å) reveals the α2β5 loop being the structurally most divergent region. In the Ubv-G08:53BP1 complex, the loop contains three evolved residues and contributes to target binding together with evolved residues from the strands β1 and β5. In spite of significant differences in details of their binding modes, Af2 and Ubv-G08 resemble each other by employing a -4 register shift of β5 to accomplish a loop extension and thereby modulating their target binding interfaces of similar sizes (supplementary figures S9e-f and i, supplementary figure S1).

### Figure S1

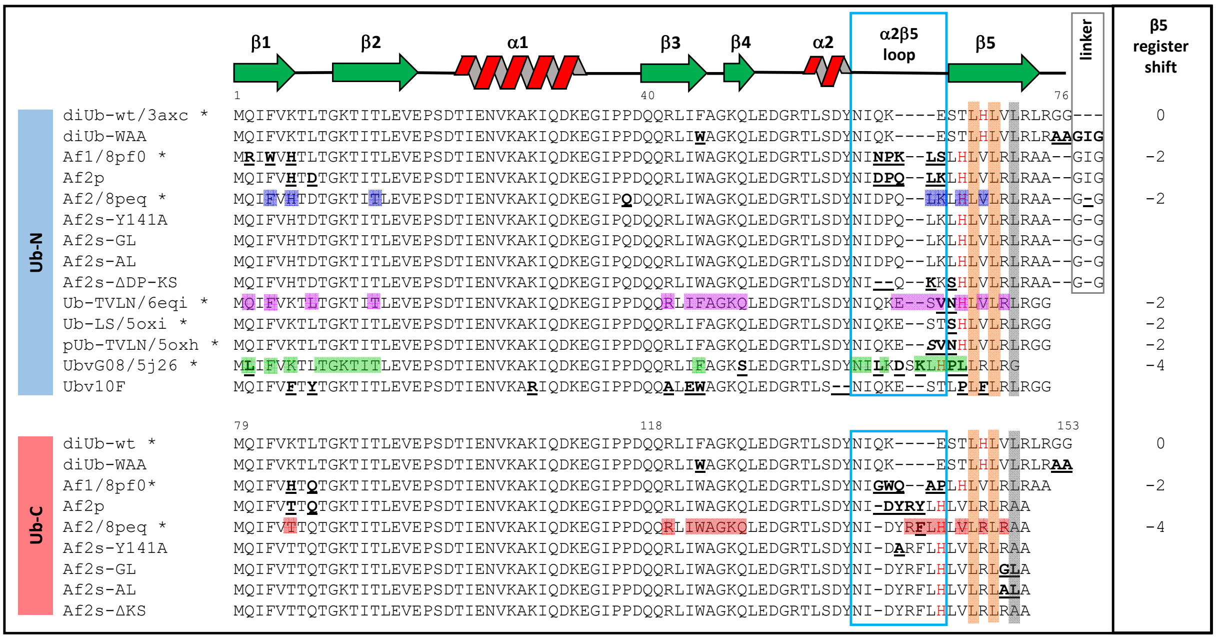

**Figure S1:** Structure-based sequence alignment of diubiquitin-based Affilin variants, diubiquitin (diUb) and selected (mono) ubiquitin variants (marked with an asterisk and PDB accession code if experimental structure is available). Upper panel: alignment of N-terminal Ub domains (or mono-Ub domains), lower panel: alignment of C-terminal Ub domains. Residue numbering and secondary structure assignment according to Af2. Sequence alterations to respective parental Affilin variant (or Ub-wt) indicated in bold and underlined (except linker residues). Residues located in the α2β5 loop are boxed in blue, linker residues in grey box, positions of Leu residues in β5 of Ub-wt shaded in orange and grey, and "marker residue" H68 of Ub-wt colored in red. Residues identified in the complex structures involved in binding are color-shaded. Register shifts in β5, if observed, given in the right column.

#### Figure S2

a

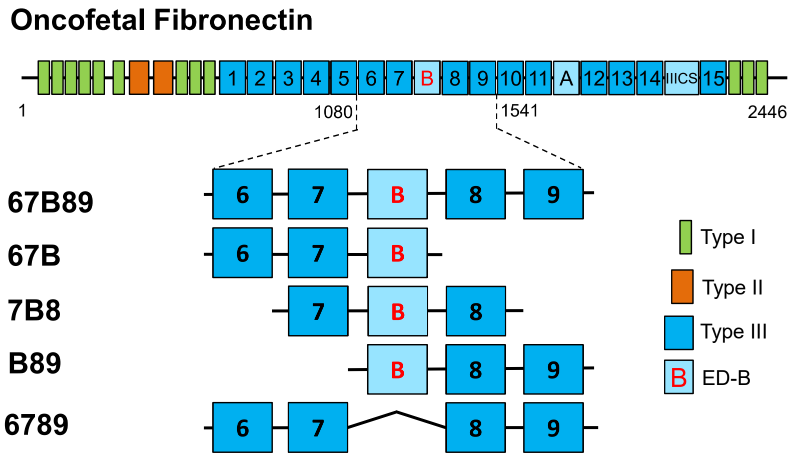

b

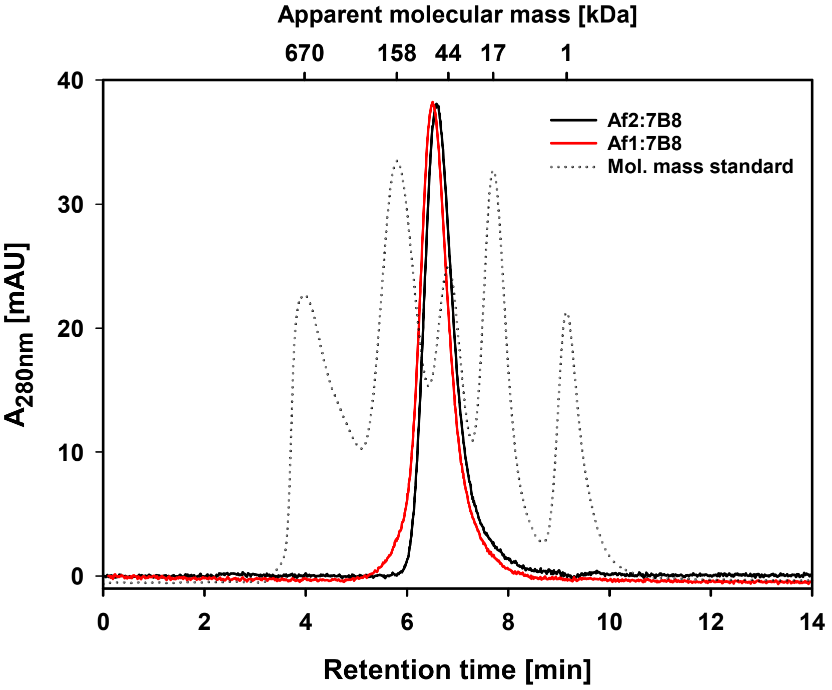

**Figure S2. (a)** Fibronectin fragments derived from oncofetal fibronectin used in this work. Based on the amino acid sequence of human fibronectin isoform 7 comprising ED-B (UniProtKB database ID: P02751, top), several truncated constructs were created for selection and maturation of binders (target 67B89, off-target 6789 lacking ED-B), binding analysis (all variants) and crystallization (7B8). The numbers represent the N- and C-terminal residues, respectively, flanking each construct, following the residue numbering in the database entry. **(b)** Size exclusion HPLC analysis of the Af2:7B8 and Af1:7B8 complexes. The preformed complexes were analyzed using a Superdex 200 5/150 GL analytical size exclusion column which had been calibrated with a HPLC gel filtration molecular weight standard consisting of Thyroglobulin (670 kDa), Gamma-globulin (158 kDa), Ovalbumin (44 kDa), Myoglobin (17 kDa) and vitamin B12 (1 kDa). Apparent molecular weights were derived from the retention times of the Af2:7B8 complex (black solid line) and the Af1:7B8 complex (red solid line), respectively, using the molecular weight standard (dotted line).

#### Figure S3

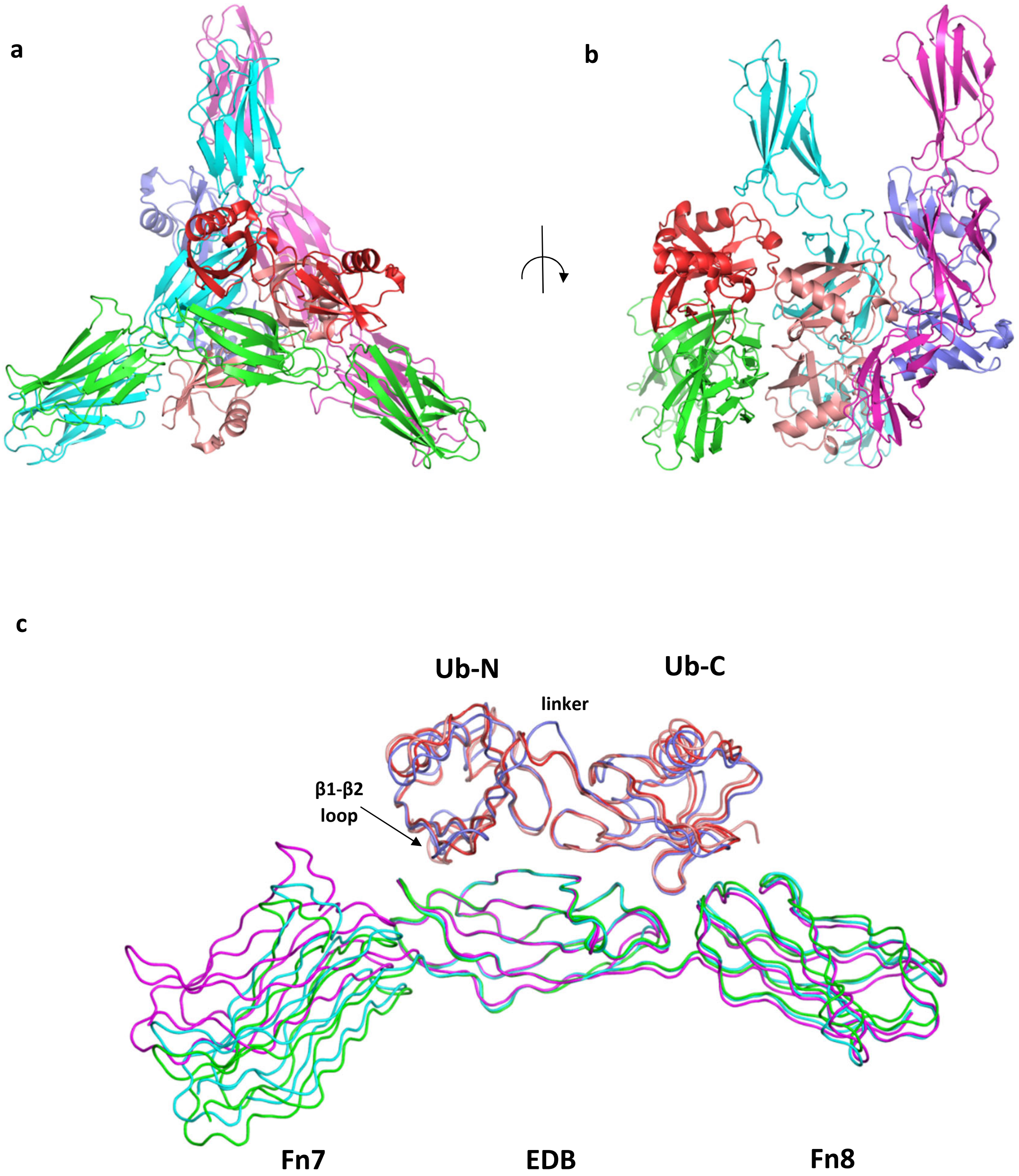

**Figure S3:** Asymmetric unit of the Af2:7B8 complex crystal structure composed by three complexes. (**a**) $3_1$-fold non-crystallographic symmetry relation of the three complexes colored by chain (7B8 : chain A – green, chain B – cyan, chain C – magenta, Af2 : chain L – red, chain J – salmon, chain M – blue) **(b)** 90° rotated view (**c**)**:** Ribbon representation of a structural comparison of the three Af2:7B8 complexes from the asymmetric unit. For superposition, only the EDB residues (1266 – 1356) of the 7B8 chains were aligned.

#### Figure S4

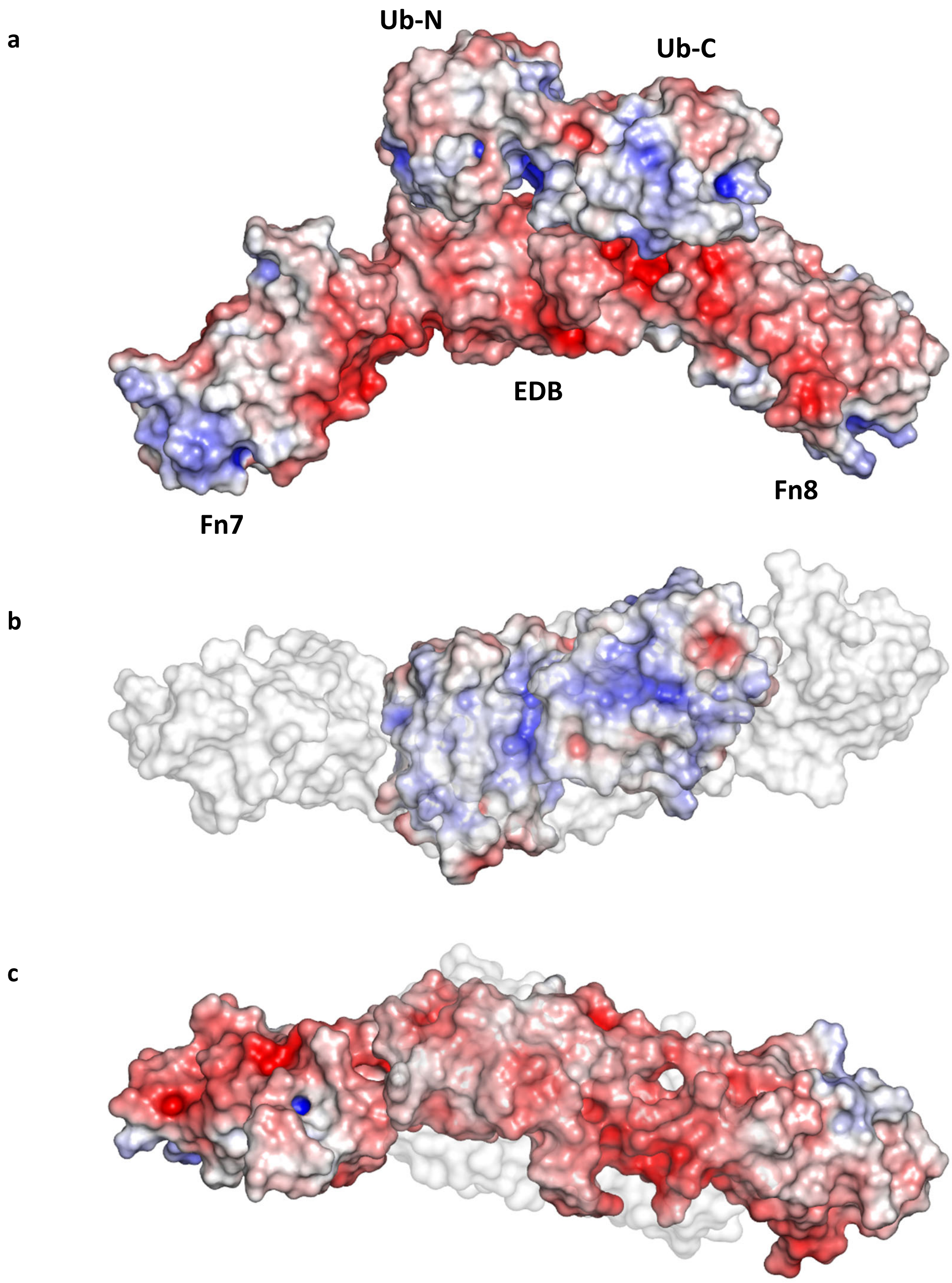

**Figure S4** Electrostatic properties of Af2 and 7B8. ±5 kT/e electrostatic potential APBS Electrostatic surface potentials were calculated using the software APBS [9] with the non-linear Poisson-Boltzmann equation contoured at 5 kT/e. Negatively and positively charged surface areas are colored in red and blue, respectively. (**a**) Side view in the orientation of figure 1d. (**b**) Rotated view on the 7B8-binding interface of Af2, the bound target 7B8 is schown as transparent white surface. (c) Rotated view on the Af2-binding face of 7B8, bound Af2 is shown as transparent white surface.

#### Figure S5

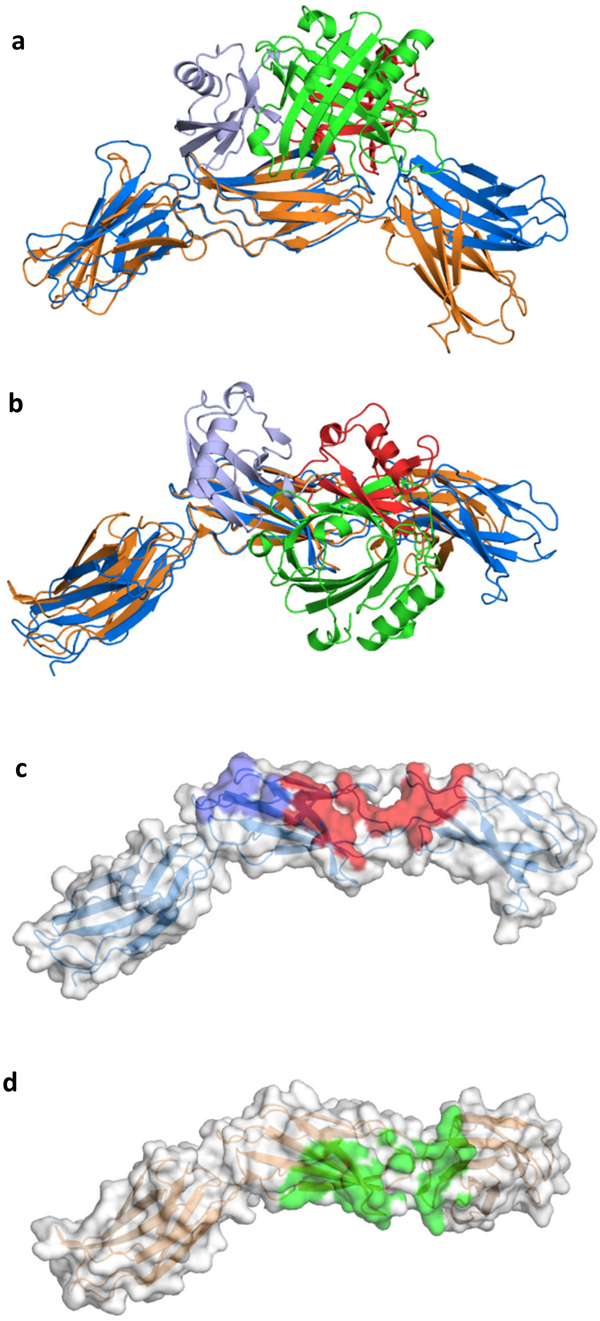

**Figure S5** Structural comparison of the Af2:7B8 complex and 7B8-bound Anticalin N7A ([10], PDB id 4GH7) (**a**) Superposition of overall structures based on alignment of the EDBs shown in the orientation of figure 1d. Af2 colored in light blue (Ub-N) and red (Ub-C), Af2-bound 7B8 colored in blue, N7A colored im green, N7A-bound 7B8 colored in orange. (**b**) 90° rotated view. (**c**) Molecular surface of Af2-bound 7B8 with mapped binding interfaces of Ub-N (blue surface) and Ub-C (red surface). (**d**) Molecular surface of N7A-bound 7B8 with mapped binding interface of N7A (green surface).

### Figure S6

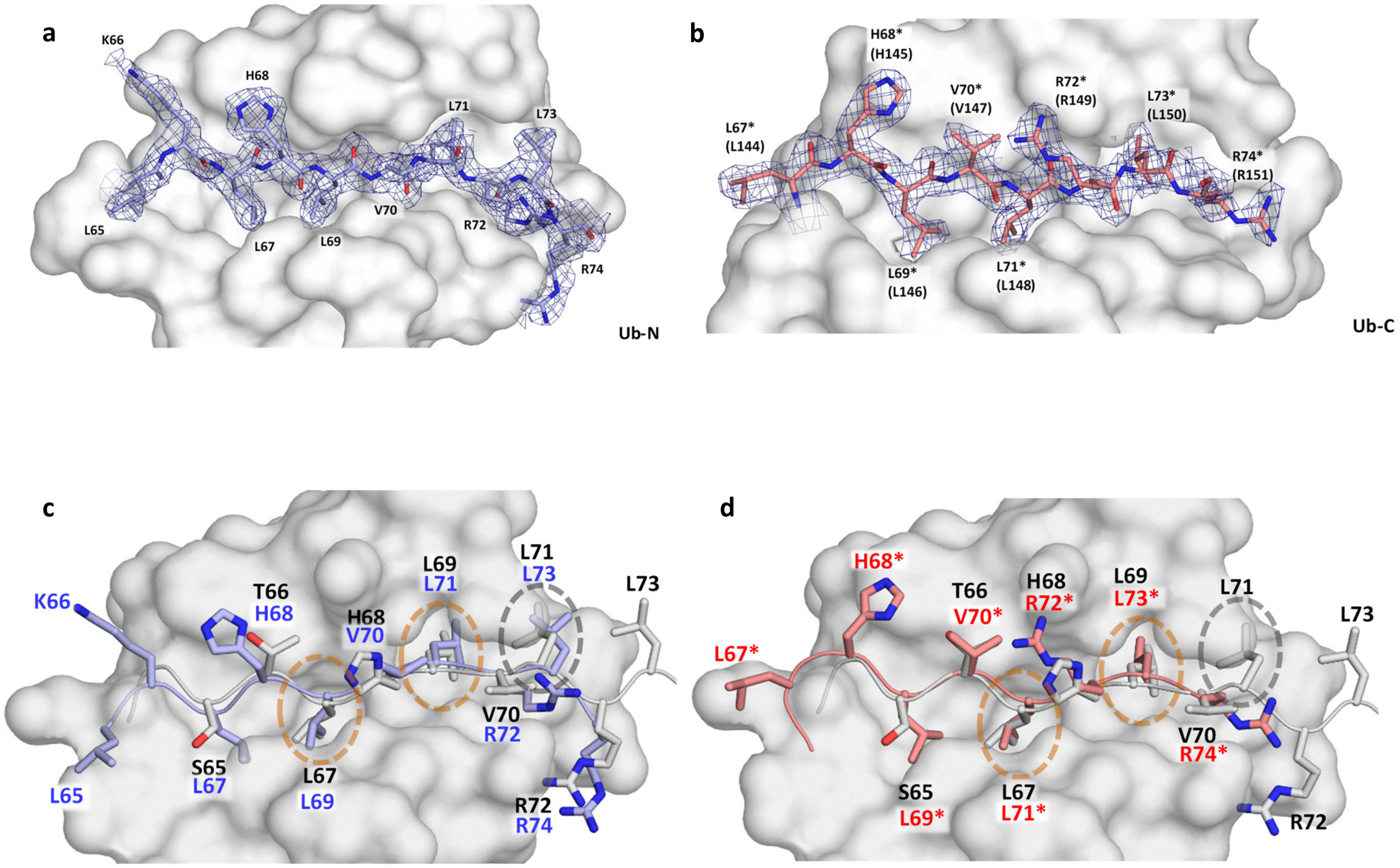

**Figure S6** (**a**) 2F$_o$-F$_c$ electron density (blue mesh) countoured at 1$\sigma$ of residues from the β5 strand of Af2 Ub-N, revealing a -2 register shift. Residues are shown as sticks, Ub-N residues (except from β5) shown as white surface. (**b**) 2F$_o$-F$_c$ electron density (blue mesh) countoured at 1$\sigma$ of residues from the β5 strand of Af2 Ub-C, revealing a -4 register shift. Residues are shown as sticks, Ub-N residues (except from β5) shown as white surface. Residue labels marked with an asterisk corresponds to the residue numbers of Ub-N. (**c**) Structural superposition of residues from the β5 strand of Af2 Ub-N (light blue sticks, blue labels) and Ub-wt ([2], PDB id 1UBQ, white sticks, black labels) depicting the -2 register shift. (**d**) Structural superposition of residues from the β5 strand of Af2 Ub-C (light red sticks, red labels) and Ub-wt (white sticks, black labels) depicting the -4 register shift. The two hydrophobic binding grooves harboring leucine residues in the non-shifted and -2/-4 register-shifted states are marked with orange circles. The grey circle marks a leucine-harboring site occupied only in the non-shifted and -2-shifted state.

#### Figure S7

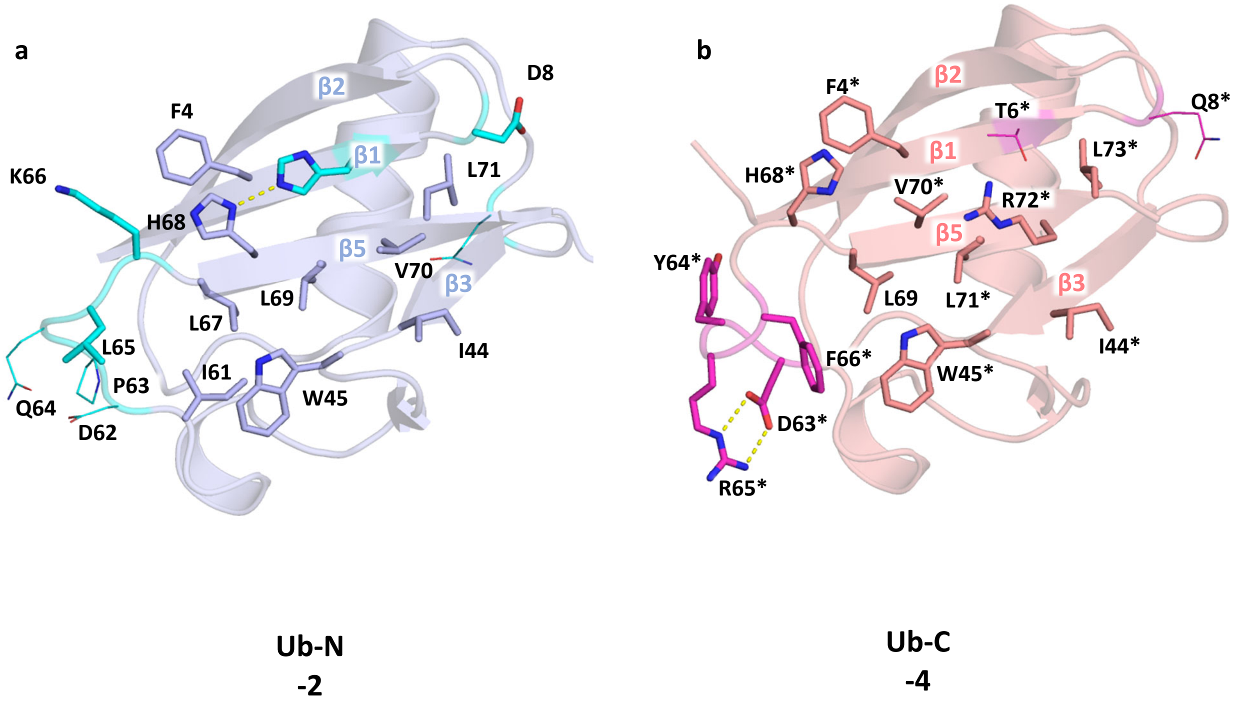

**Figure S7** Intramolecular interactions stabilizing the register-shifted states observed in Af2. (**a**) Residues involved in stabilizing the -2 shift in Ub-N shown as sticks, evolved amino acids colored in cyan, shown as lines if not involved in intramolecular stabilization of the shift. (**b**) Residues involved in stabilizing the -4 register shift in Ub-C shown as sticks, evolved amino acids colored in magenta, shown as lines if not involved in intramolecular stabilisation of the shift. Residue labels marked with an asterisk corresponds to the residue numbering of Ub-N.

#### Figure S8

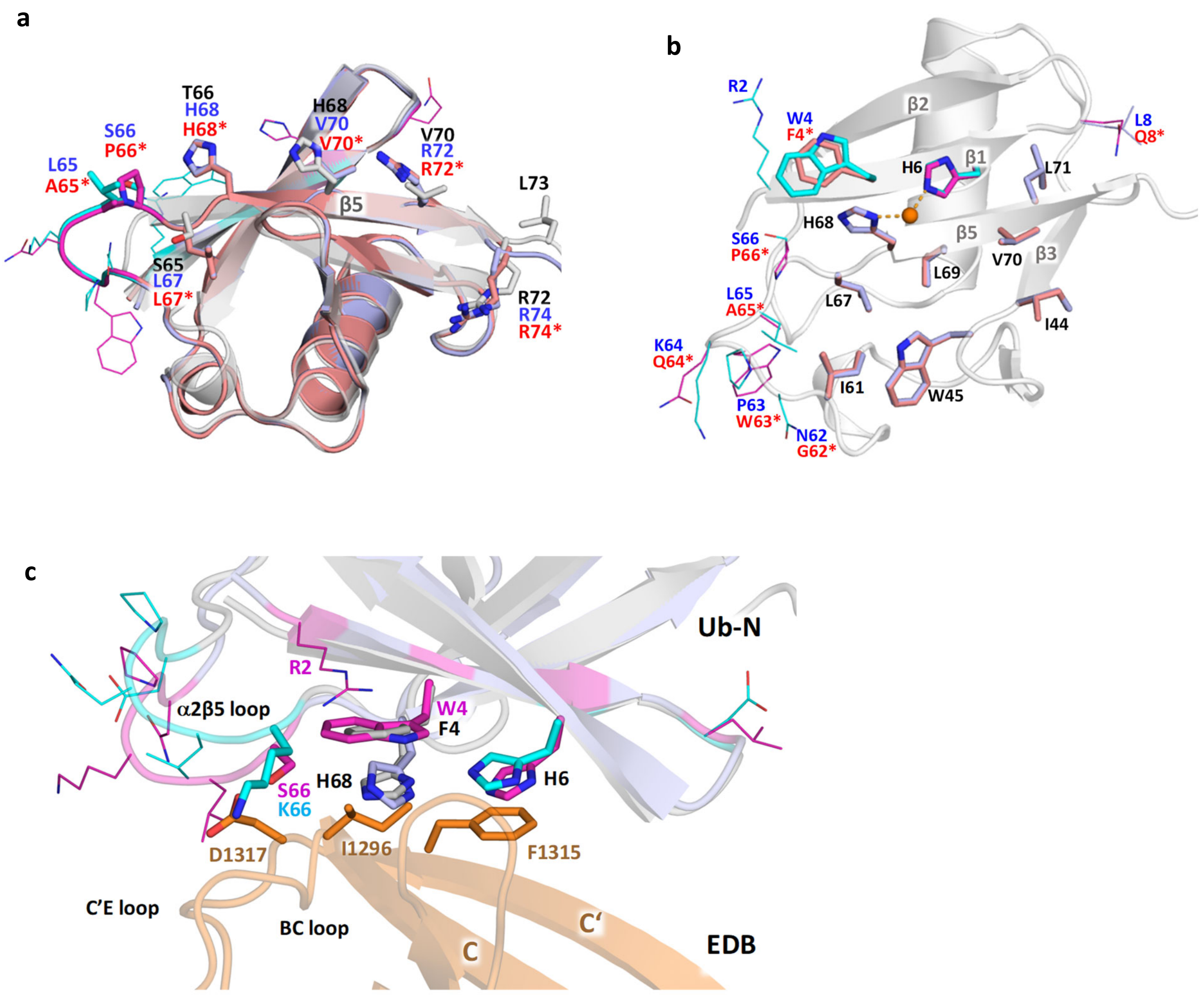

**Figure S8** The -2 register shifts observed in the crystal structure of unbound Af1. **(a)** Structural superposition of Ub-N (light blue, blue labels) and Ub-C (light red, red labels) of Af1, both displaying a -2 register-shift in β5 vs. Ub-wt (PDB id 1UBQ, white, black labels). Residues of β5 shown as sticks. Evolved Af1 residues in Ub-N shown in cyan, evolved residues in Ub-C shown in magenta. Evolved residues outside β5 shown as lines. (**b**) Superimposed residues of Ub-N and Ub-C involved intramolecular stabilization of the -2 shift in Af1, shown as sticks, evolved amino acids colored in cyan (Ub-N, blue labels) and magenta (Ub-C, red labels). Residues shown as lines if not involved in intramolecular stabilization of the shift. Residue labels marked with an asterisk corresponds to the residue numbering of Ub-N, residues are identical in Ub-N and Ub-C are labelled in black. H68 and H6 of Ub-N and Ub-C coordinate a copper ion (orange sphere and dashes) present in the crystallisation solution. **(c)** Structural superposition of the Ub-N domains of target-bound Af2 (Ub-N colored in grey, evolved side chains in cyan, EDB in orange) and unbound Af1 (colored in light blue, evolved side chains in magenta). Residues of the binding interface I in the Af2:7B8 complex and corresponding residues in Af1 shown as sticks, other evolved residues shown as lines.

**Figure S9**

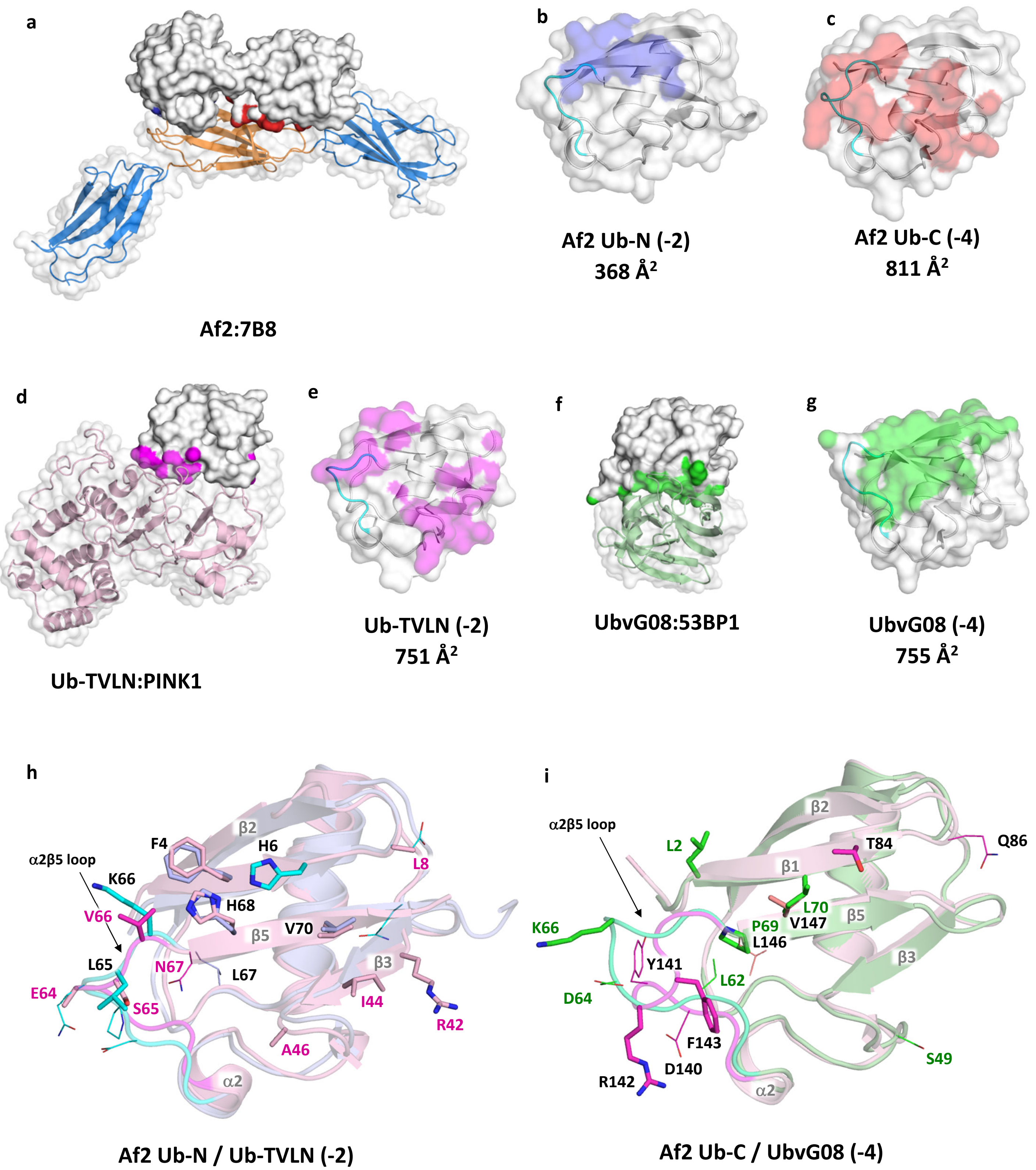
a
Af2:7B8
b
Af2 Ub-N (-2)
368 Å²
c
Af2 Ub-C (-4)
811 Å²
d
Ub-TVLN:PINK1
e
Ub-TVLN (-2)
751 Å²
f
UbvG08:53BP1
g
UbvG08 (-4)
755 Å²
h
α2β5 loop
F4
β2
H6
L8
K66
H68
V66
β5
V70
L65
N67
L67
β3
E64
I44
R42
S65
A46
α2
Af2 Ub-N / Ub-TVLN (-2)
i
α2β5 loop
β2
L2
T84
Q86
β1
L70
K66
P69
V147
β5
L146
Y141
L62
β3
D64
F143
S49
D140
R142
α2
Af2 Ub-C / UbvG08 (-4)

**Figure S9** Structural comparison of the Af2:7B8 complex with other Ub complexes exhibiting register shifts in the β5 strand. (**a**) Molecular surface representation of the Af2:7B8 complex. Af2 shown as white solid surface with mapped 7B8 binding interfaces of Ub-N (blue surface) and Ub-C (red surface). 7B8 shown as transparent surface, EDB colored orange, Fn7 and Fn9 colored blue. (**b**) The 7B8-binding interface of Af2 Ub-N, mapped in blue on white domain surface, α2β5 loop colored in cyan. (**c**) 7B8-binding interface of Af2 Ub-C, mapped in red on domain surface, α2β5 loop colored in cyan. (**d**) Molecular surface representation of the Ub-TVLN:PINK1 complex (PDB id 6eqi). Ub-TVLN shown as white solid surface with mapped PINK1 binding interfaces of the Ub domain (magenta surface). PINK1 shown as transparent surface, overall structure colored in light pink. (**e**) PINK1-binding interface of Ub-TVLN, mapped in pink on Ub domain surface, α2β5 loop colored in cyan. (**f**) Molecular surface representation of the UbvG08:53BP1 complex (PDB id 5j26). UbvG08 shown as white solid surface with mapped 53BP1 binding interfaces of the Ub domain (green surface). 53BP1 shown as transparent surface, overall structure colored in light green. (**g**) 53BP1-binding interface of UbvG08, mapped in green on Ub domain surface, α2β5 loop colored in cyan. (**h**) Structural superposition of Af2 Ub-N (light blue, black labels, evolved residues and α2β5 loop colored in cyan) and Ub-TVLN (light red, magenta labels, α2β5 loop residues colored in magenta), both displaying a -2 register-shift in β5. Residues involved in binding shown as sticks. Altered residues not contributing to binding shown as lines. (**i**) Structural superposition of Af2 Ub-C (light red, black labels, evolved residues and α2β5 loop colored in magenta) and UbvG08 (light green, green labels, α2β5 loop residues colored in cyan), both displaying a -4 register-shift in β5. Residues involved in binding shown as sticks. Evolved residues not contributing to binding shown as lines.

#### Table S1: Statistics of data collection and structure refinement

| Dataset (PDB accession) | Af1 (8PF0) | Af2:7B8 complex (8PEQ) |
|---|---|---|
| X-ray source | BESSY BL14.2 | BESSY BL14.1 |
| wavelength [Å] | 0.9184 | 0.9184 |
| Detector | CCD MX225 | PILATUS 6M |
| space group | $P4_1$ | C2 |
| Cell parameter | | |
| a,b,c [Å] | 62.77, 62.77, 67.84 | 168.67, 105.86, 78.99 |
| α,β,γ [°] | 90.00, 90.00, 90.00 | 90.00, 93.52, 90.00 |
| Resolution [Å] | 30.0-2.2 | 43.0-2.3 |
| | (2.3-2.2) | (2.4-2.3) |
| completeness [%] | 99.5 (97.6) [1] | 96.8 (78.7) [2] |
| total reflections | 99386 | 210486 |
| unique reflections | 26176 (3189) [1] | 59628 (5777) [2] |
| multiplicity | 3.8 (3.8) [1] | 3.5 (3.3) [2] |
| $R_{merge}$ | 3.1 (38.7) [1] | 5.8 (39.9) [2] |
| I/σ(I) | 22.9 (3.9) [1] | 16.5 (3.1) [2] |
| $CC_{1/2}$ | 99.9 (91.0) [1] | 99.8 (83.9) [2] |
| Wilson B-factor | 58.6 | 41.9 |
| **Structure refinement** | | |
| molecules per | 1 Af1 | 3 Af2 |
| asymmetric unit | | 3 7B8 |
| R values [%] | | |
| $R_{work}$ | 19.3 | 19.8 |
| $R_{free}$ | 22.6 | 25.1 |
| number of atoms | | |
| protein | 1223 | 9940 |
| $Cu^{2+}$ ions | 4 | - |
| buffer components | 10 | 10 |
| solvent | 78 | 536 |
| average B factor [$Å^2$] | 55.6 | 39.8 |
| Rmsd | | |
| bond lengths [Å] | 0.007 | 0.008 |
| bond angles [°] | 1.1 | 1.0 |
| Ramachandran [%] | | |
| favored | 98.0 | 97.9 |
| allowed | 2.0 | 2.0 |
| outlier | 0.0 | 0.1 |
| Molprobity clashscore | 2.4 | 4.5 |

Values for highest resolution shell are given in parentheses.

[1] Friedel pairs treated as independent reflections.

[2] Friedel pairs merged.
